# Re-evaluating the actin-dependence of spectraplakin functions during axon growth and maintenance

**DOI:** 10.1101/2021.11.21.469398

**Authors:** Yue Qu, Juliana Alves-Silva, Kriti Gupta, Ines Hahn, Jill Parkin, Natalia Sánchez-Soriano, Andreas Prokop

## Abstract

Axons are the long and slender processes of neurons constituting the biological cables that wire the nervous system. The growth and maintenance of axons require bundles of microtubules that extend through their entire length. Understanding microtubule regulation is therefore an essential aspect of axon biology. Key regulators of neuronal microtubules are the spectraplakins, a well-conserved family of cytoskeletal cross-linkers that underlie neuropathies in mouse and humans. Spectraplakin deficiency in mouse or *Drosophila* causes severe decay of microtubule bundles and axon growth inhibition. The underlying mechanisms are best understood for *Drosophila* Short stop (Shot) and believed to involve cytoskeletal cross-linkage: the N-terminal calponin homology (CH) domains bind to F-actin, and the C-terminus to microtubules and Eb1. Here we have gained new understanding by showing that the F-actin interaction must be finely balanced: altering the properties of F-actin networks or deleting/exchanging Shot’s CH domains induces changes in Shot function - with a Lifeact-containing Shot variant causing remarkable remodelling of neuronal microtubules. In addition to actin-MT cross-linkage, we find strong indications that Shot executes redundant MT bundle-promoting roles that are F-actin-independent. We argue that these likely involve the neuronal Shot-PH isoform, which is characterised by a large, unexplored central plakin repeat region (PRR). Work on PRRs might therefore pave the way towards important new mechanisms of axon biology and architecture that might similarly apply to central PRRs in mammalian spectraplakins.

## Introduction

Axons are the slender, up-to-two-meter-long processes of neurons that form the biological cables wiring our bodies (Prokop, 2020). Their *de novo* formation during development, regeneration or brain plasticity is implemented at growth cones (GCs), the amoeboid tips of extending axons (Harrison, 1910; Ramón y Cajal, 1890). GCs navigate by sensing spatiotemporally patterned chemical and mechanical cues along their paths which are translated into orchestrated morphogenetic changes leading to axon extension (Franze et al., 2013; Sanes et al., 2019; Tessier-Lavigne and Goodman, 1996).

These morphogenetic changes are mediated by the cytoskeleton, in particular, actin and microtubules (MTs; Dent et al., 2011; Lowery and van Vactor, 2009; Prokop et al., 2013; Tanaka and Sabry, 1995): F-actin in the GC periphery is required for explorative protrusive activity and mechano-sensing and will eventually mediate the directional stabilisation of MTs which will, in turn, implement the actual growth events (e.g. Buck and Zheng, 2002; Geraldo et al., 2008; Lee and Suter, 2008; Qu et al., 2019; Suter and Forscher, 2001). If MTs in GCs arrange into bundled loops or spools, they seem to suppress such interactions in the periphery and slow down axon growth (Dent et al., 1999).

The MTs of GCs originate from the MT bundles of the axon shaft. These bundles run all along axons and serve as the essential highways for axonal transport (Prokop, 2020). They must therefore be maintained throughout an organism’s lifetime involving active repair and turn-over (Hahn et al., 2019; Prokop, 2021). These bundles can also drive axon elongation through so-called intercalative or stretch growth (Bray, 1984; Lamoureux et al., 2010; Smith, 2009; Zheng et al., 1991). For this, axons display forward drift of MT bundles (Miller and Sheetz, 2006; Roossien et al., 2013) or MT sliding forces (Lu et al., 2015; Winding et al., 2016). Like in GCs, the MT bundle regulation in axon shafts requires actin-MT interactions required for their parallel arrangement and to uphold MT numbers (Alves-Silva et al., 2012; Datar et al., 2019; Krieg et al., 2017; Qu et al., 2017).

Numerous mechanisms have been described that mediate actin-MT interaction (Dogterom and Koenderink, 2019; Kundu et al., 2021; Mohan and John, 2015). In axons, very prominent mediators are the spectraplakins, an evolutionarily well-conserved family of multi-domain cytoskeletal linker proteins (Fig.1A; Voelzmann et al., 2017). Of these, dystonin was discovered in a mouse model of sensory neuropathy, later shown to involve severe MT bundle deterioration and be linked to human HSAN6 (hereditary sensory and autonomic neuropathy: OMIM #614653; Duchen et al., 1964; Edvardson et al., 2012; Eyer et al., 1998). Its mammalian paralogue ACF7/MACF1 was discovered as an actin-MT cross-linker (Byers et al., 1995; Leung et al., 1999), later shown to be involved in neuronal development (Goryunov et al., 2010; Ka et al., 2014; Ka and Kim, 2015; Sánchez-Soriano et al., 2009) and linked to lissencephaly (OMIM #618325). As detailed elsewhere (Voelzmann et al., 2017), spectraplakins can act as actin-MT cross-linkers: they bind F-actin via a tandem of N-terminal calponin homology domains (CH domains) and associate with MTs through their C-terminus; this C-terminus harbours a GRD (Gas2-related domain) which also stabilises MTs against depolymerisation, and a positively charged unstructured Ctail which also binds to Eb1 (Fig.1A; Alves-Silva et al., 2012; Goriounov et al., 2003; Honnappa et al., 2009; Lee and Kolodziej, 2002).

**Fig.1.**
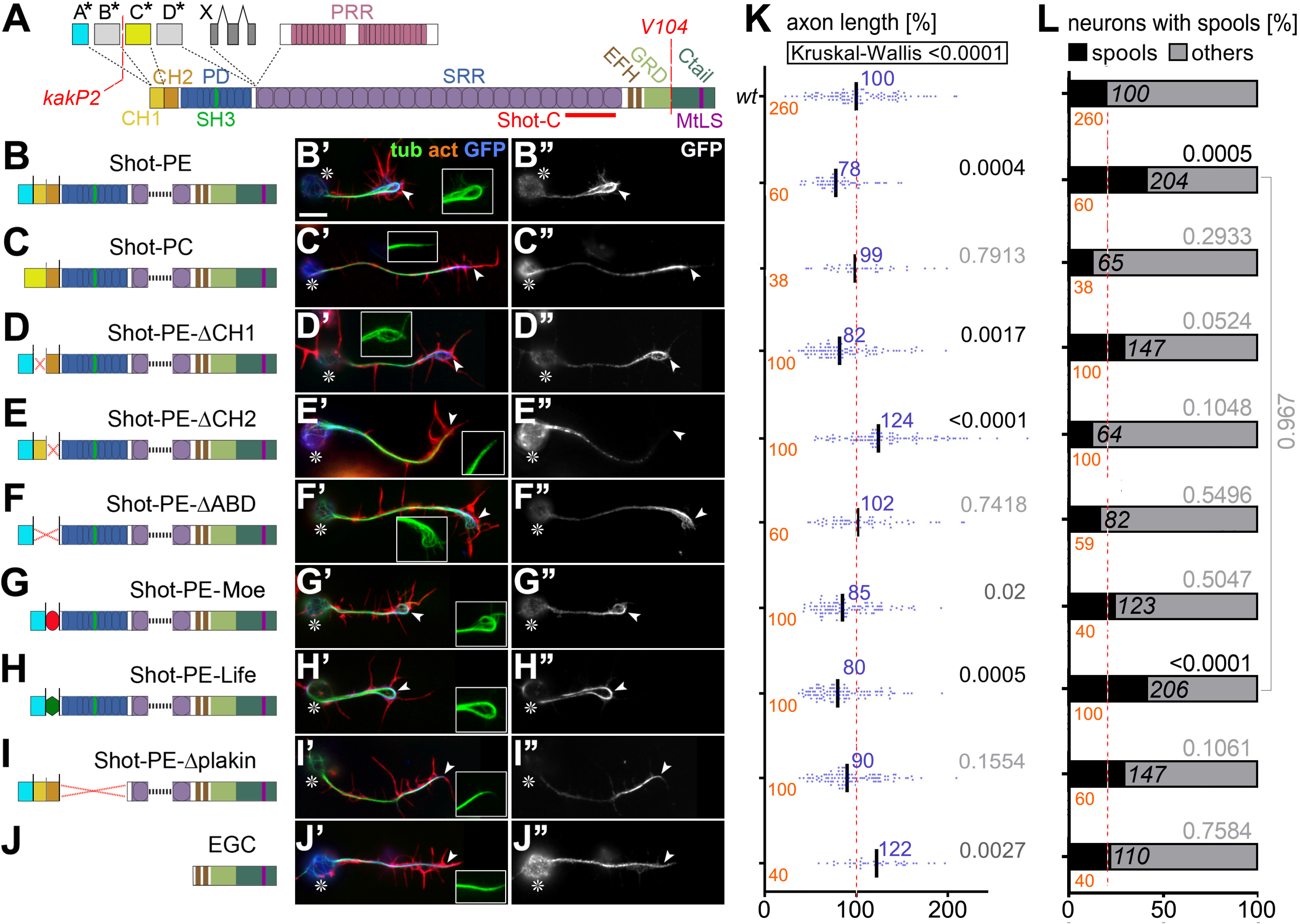
Different Shot constructs and their localisation. **A**) Illustration of different Shot isoforms as a function of different start sites (A*-D*) and splice-in of different exons (X, PRR); different domains and motifs are colour-coded (CH, calponin homology; PD, plakin domain; PRR, plakin repeat region; SRR, spectrin repeat region; EFH, EF-hand; GRD, Gas2-related domain; MtLS, MT tip localization sequence which forms the Eb1-binding motifs); positions of the epitope used to generate the Shot-C antibody (Strumpf and Volk, 1998), the *kakP2* P-element insertion (blocking the A* and B* start sites) and the break-point of the *V104* inversion (deleting the Ctail) are indicated in red. **B-J**) Different UAS-constructs expressing modified Shot versions. **B’-J’**) Primary neurons at 6-8 HIV cultured on glass which express the respective constructs on the left and are stained for actin (red), tubulin (green) and GFP (blue). **B’’-J’’**) GFP channel shown in grayscale. In all images, asterisks indicate cell bodies, arrow heads the axon tips; scale bar in A represents 10 µm in all images. **K, L**) Graphs display the distribution of axon length phenotypes (K) and frequency of spools in neuronal GCs (L) taken from neuron populations expressing the same constructs as displayed in B-J’’. Number of neurons analysed are shown in orange, median values in blue (K only), black numbers within columns in L indicate the percentage of neurons with spool-containing GCs; black/grey numbers on the right of each plot/bar indicate the P-values obtained via Mann–Whitney Rank Sum Tests in K (Kruskall-Wallis ANOVA test results shown above) and Chi-square tests in L. Data were normalised to wild-type controls performed in parallel to all experiments (red dashed lines).

The *Drosophila* spectraplakin Short stop (Shot) is a close orthologue of dystonin and ACF7/MACF1. In neurons, Shot is required for axon and dendrite growth, neuronal polarity, axonal compartmentalisation, synapse formation and axonal MT bundle maintenance (Lee et al., 2000; Prokop et al., 1998; Reuter et al., 2003). In Shot-deficient neurons, MT bundles in axon shafts and GCs frequently disintegrate into disorganised, curled, criss-crossing arrangements (from now on referred to as MT curling). This dramatic MT phenotype can be rescued when reinstating actin-MT cross-linking activity of Shot, through a mechanism where Shot guides the extension of polymerising MTs along the axonal cortex into parallel bundles (Alves-Silva et al., 2012; Hahn et al., 2021; Sánchez-Soriano et al., 2010). Whereas the necessary C-terminal interaction of Shot with MTs is quite well described, we have little knowledge of the N-terminal interaction with neuronal F-actin networks, especially when considering that these can be of very different nature: presenting as sparse cortical F-actin rings in the axon shaft (Leterrier et al., 2017) or dense F-actin networks in GCs (Dent et al., 2011).

Here, we have gained new understanding of Shot’s F-actin interaction. Firstly, we show that Shot function does not simply depend on F-actin: it rather appears to involve a well-balanced interplay of low-affinity CH domains with F-actin networks, where any changes can trigger alterations in Shot’s functional output; this phenomenon is relevant for axon growth-regulating MT spool formation in GCs. In the axon shaft, Shot acts as an F-actin/MT/Eb1 cross-linker in MT bundle maintenance. In addition, we provide strong indications that Shot performs actin-independent bundle-maintaining functions acting redundantly to F-actin/MT/Eb1 cross-linkage. We argue these functions to be mediated by the Shot-PH isoform characterised by an evolutionarily conserved plakin repeat region (PRR) that is functionally unexplored (Hahn et al., 2016; Röper and Brown, 2003; Voelzmann et al., 2017) and might therefore hold the key to uncharted mechanisms of axon biology and architecture.

## Results

### Roles of Shot’s actin-binding domain in gain-of-function experiments

To assess F-actin dependency of Shot function, we first took a gain-of-function (GOF) approach. For this, we targeted the expression of transgenic Shot constructs to primary *Drosophila* neurons and analysed them at 6 hours *in vitro* (HIV) for two phenotypes: we quantified the length of axons and the number of neurons showing bundled loops referred to as ‘spools’ (Fig.1B’,G’,H’) - as opposed to ‘pointed’ (Fig.1C’,D’,I’,J’) or ‘disorganised’ (Fig.1F’; Sánchez-Soriano et al., 2010; Teng et al., 2001). Neuronal expression of Shot-PE::GFP (a GFP-tagged version of the best-studied Shot isoform; Hahn et al., 2016; Fig.1A,B), caused a reduction in axon length to ∼80% and doubled the number of MT spools in growth cones (GCs) when compared to wild-type controls (Figs.1B’,L). In contrast, Shot-PC::GFP (another natural isoform which lacks CH1; Figs.1A,C), failed to induce either of these phenotypes; instead it showed a trend to suppress spool numbers below control levels (Figs.1C’,L), as similarly observed in previous studies (Sánchez-Soriano et al., 2010). The finding suggests that an interaction with F-actin is essential for spool formation, since lack of CH1 in the Shot-PC isoform (Fig.1C) eliminates F-actin interaction (concluded from localisation and binding studies; Lee and Kolodziej, 2002). Accordingly, spool induction can also be suppressed when depleting F-actin with the drug latrunculin A (LatA; Fig.2B,D; Sánchez-Soriano et al., 2010).

**Fig.2.**
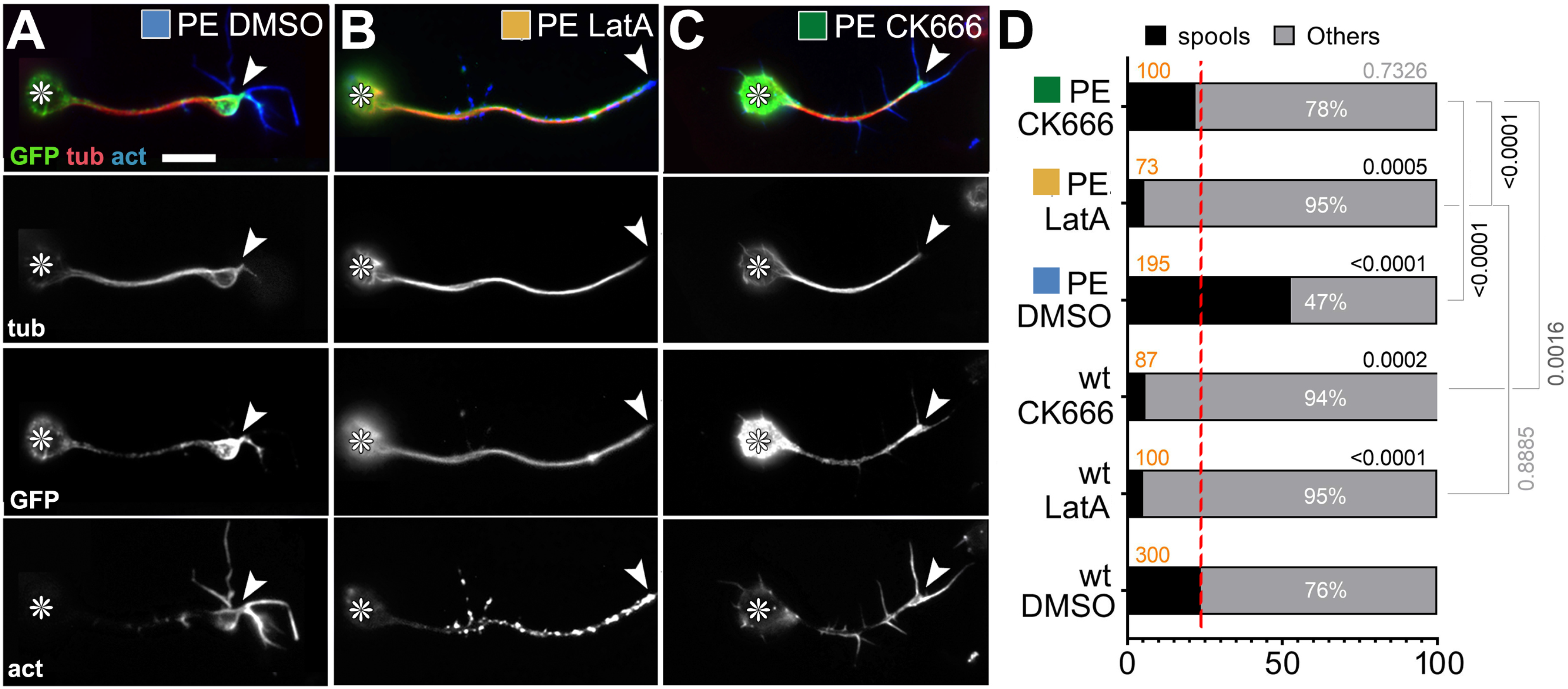
Impact of drug-induced F-actin inhibition on Shot-PE function. **A-C**) Primary neurons at 6-8 HIV on glass treated with DMSO (control), LatA or CK666 as indicated and stained for GFP (green), tubulin (red) and actin (blue); grayscale images below show single channels as indicated; asterisks indicate cell bodies, arrowheads the tips of axons; scale bar in A represents 10 µm in all images. D) Frequency of neurons with GCs that contain spools (examples of neurons in A-C are assigned to their respective data columns via colour-coded squares); orange numbers indicate the sample numbers (number of neurons analysed), black numbers within columns the percentage of neurons with GCs that contain spools; numbers on the right of each graph indicate the P-values obtained via Chi^2^ tests. Data were normalised to wild-type controls performed in parallel to all experiments (dashed red line).

Shot-PE and Shot-PC not only differ in the presence/absence of CH1, they also display different lead sequences that flank CH domains N-terminally (blue A* *vs.* yellow C* in Fig.1A-C; Hahn et al., 2016). Both lead sequences lack any informative homologies or motifs but may still be functionally relevant, for example by having different modifying impacts on CH domain functions (Yin et al., 2020). Therefore, we generated Shot-PE-ΔABD::GFP, a Shot-PE variant containing the A* lead sequence but lacking both CH domains (Fig.1D-F). The phenotypes observed upon Shot-PE-ΔABD::GFP expression were almost identical to those of Shot-PC (Fig.1F’,K,L), corroborating former claims that the actin-binding capability of Shot-PC is negligible (Lee and Kolodziej, 2002).

Surprising results were obtained when deleting single CH domains in the Shot-PE context. Previous work suggested that CH1 is the main actin-binding domain of the tandem (Korenbaum and Rivero, 2002; Lee and Kolodziej, 2002; Sjöblom et al., 2008; Yin et al., 2020), and we expected therefore that Shot-PE-ΔCH2 would have modest actin-binding hence spool-inducing capability, whereas Shot-PE-ΔCH1 would be similar to Shot-PC or Shot-ΔABD. However, we found the opposite: When deleting the functionally less prominent CH2, we found robust axon elongation to ∼120% and failure to induce extra spools, i.e. a phenotype suggesting complete loss of actin-binding properties although CH1 was present (Fig.1E,K,L). In contrast, Shot-PE-ΔCH1 expression had a trend towards extra spool formation and shorter axons, suggesting modest actin-binding properties although CH1 was absent. Shot-PE-ΔCH1 resembles Shot-PC in that it lacks the CH1 domain, but it contains the A* lead sequence instead of C* (Figs.1C *vs.* D). Our results might therefore hint at potential regulatory roles of the N-terminal lead sequences: for example, the C* lead sequence of Shot-PC, but not the A* sequence of Shot-PE, might inhibit residual actin affinities of CH2, thus explaining why Shot-RE-ΔCH1 appears to display more activity than Shot-RC and Shot-RE-ΔABD.

### F-actin is required for Shot construct localisation

To gain more understanding of these phenotypes, we performed localisation studies. Shot-PE::GFP is strongly enriched at the distal end of axons, mostly at the actin-enriched growth cones (GCs); this is consistent with its spool-inducing activity (Fig.1B’’). Also, Shot-PC::GFP and Shot-PE-ΔABD::GFP are distally enriched in axons (Fig.1C’’,F’’), suggesting that their inability to induce spools is not due to their physical absence but rather their functional impairment.

Also Shot-PE-ΔCH1::GFP is enriched in distal axon segments (Fig.1D’’). This localisation is consistent with its spool-inducing tendencies which might be mediated by residual F-actin affinity of its CH2 domain (see above). In contrast, the Shot-PE-ΔCH2::GFP construct is retained at or actively localises to proximal axon segments (Fig.1E’) which is consistent with the absence of its spool-inducing activity (Fig.1L).

It is surprising that even Shot constructs lacking their CH domains localise distally at F-actin-rich GCs, although this distal localisation was nevertheless F-actin-dependent: LatA treatment abolished the distal accumulation of Shot-PE::GFP (’GFP’ in Fig.2B) as was similarly observed with the F-actin-inhibiting drug cytochalasin D (CytoD; Fig.S1B).

C-terminal domains of Shot seem not to be involved since the GFP-tagged C-terminus (Shot-EGC::GFP; comprising EF-hand motifs and the MT-binding GDR and Ctail; Fig.1J) localises homogeneously along axonal MTs, and does not induce extra spools or axon shortening (Fig.1J-L; Alves-Silva et al., 2012). Instead, we focussed on the N-terminal plakin domain, because Shot-PE-Δplakin::GFP had been reported to display transient localisation defects in developing embryonic motor nerves (Bottenberg et al., 2009). However, like most other constructs, Shot-PE-Δplakin::GFP displayed distal localisation in primary neurons (Fig.1I’’), but it failed to induce robust spool formation or axon shortening (Fig.1K,L; consistent with its partial deficits in supporting axon growth *in vivo*; Bottenberg et al., 2009).

Taken together, our data suggest complex regulations at the N-terminus. We propose that two domains can mediate F-actin association: CH domains through direct binding, and the plakin domain (which contains a SRC Homology 3 motif of protein interaction; ‘SH3’ in Fig.1A) through association with independent factors that are localised at GCs through F-actin (e.g. transmembrane proteins; see Discussion). In this scenario, distal localisation of Shot could be mediated by either the CH domains or the plakin domain alone, but its spool-inducing function would depend on both domains in parallel; this would explain why single deletion of either the plakin or the CH domains abolishes Shot’s spool-inducing activity but not its localisation.

### Qualitative or quantitative changes of F-actin interaction influence Shot’s MT-regulating roles

As explained above, we propose that Shot interacts with F-actin networks through both the plakin and CH domains. This raises the question of whether Shot uses F-actin as a mere anchor or whether its function is influenced by changes in the quantity and quality of F-actin networks. To address this, we first introduced quantitative and qualitative changes to F-actin networks by manipulating actin nucleation, i.e. the process of seeding new actin filaments.

In *Drosophila* primary neurons, nucleation is performed primarily by the formin DAAM and the Arp2/3 complex (Gonçalves-Pimentel et al., 2011; Prokop et al., 2011); of these, Arp2/3 is expected to contribute branched networks that are qualitatively different from those nucleated by formins (Blanchoin et al., 2014). Arp2/3-mediated actin nucleation can be specifically inhibited by CK666 (Hetrick et al., 2013). When applying 100 nM CK666 for 2 hrs, we observed a reduction in filopodia numbers to 72±5% (P_Mann-Whitney_<0.001, n=80), indicating successful Arp2/3 inhibition and a reduction in F-actin abundance (Gonçalves-Pimentel et al., 2011). Under these conditions, Shot-PE::GFP was still recruited to the distal axon, but its spool-inducing activity was strongly suppressed (Fig.2C,D). This finding supports our hypothesis that quantitative and/or qualitative changes of F-actin networks impact MT regulatory roles of Shot.

To further challenge this notion, we decided to exchange the two CH domains of Shot for conceptually different actin-binding domains taken from other proteins. For this, we chose the 17 residue actin-binding motif Lifeact (Life) from the *Saccharomyces cerevisiae* protein Abp140 (Riedl et al., 2008), and the C-ERMAD domain of Moesin (Moe; Kiehart et al., 2000; Millard and Martin, 2008). When extrapolating from binding studies reported for CH domains of α-actinin (closely related to those of Shot; Fig.S2), we expected that Shot’s CH domains bind F-actin modestly, whereas Life should bind F-actin more robustly in a phalloidin-like manner (Lemieux et al., 2014). In contrast, Ezrin’s actin-binding domain (closely related to Moe; Fritzsche et al., 2013; Fritzsche et al., 2014) was shown to dissociate even faster from F-actin than α-actinin’s CH domains, consistent with observations that full-length Moesin does not strongly co-localise with F-actin in embryonic chick neurons or PC12 cells (Amieva and Furthmayr, 1995; Marsick et al., 2012). We, therefore, predicted a gradual impact of the different actin-binding domains on Shot localisation and/or function in the hierarchical sequence Life > Shot CH1+2 ≥ Moe.

We first analysed the localisation of the different actin-binding domains fused to the N-terminal lead sequence of Shot-PE (GFP::A*::CH1+2, GFP::A*::Life, GFP::A*::Moe; Fig.S3) by transfecting them into *Drosophila* primary neurons. Like GFP controls, also GFP::A*::CH1+2 and GFP::A*::Moe were distributed fairly homogeneously throughout entire neurons, consistent with their expected low affinity for F-actin (Fig.S3A-C). In contrast, GFP::A*::Life showed the expected robust, phalloidin-like staining (Fig. S3D). None of the three fusion constructs caused any obvious MT phenotypes (Fig.S3E).

We next replaced both CH domains in Shot-PE::GFP with Life or Moe (Fig.1G,H) and generated transgenic flies using the same genomic landing site as utilised for other transgenic constructs in this study (see Methods); this makes sure that the expression strength was comparable between constructs (Bischof et al., 2007). When targeted to primary neurons, Shot-PE-Moe::GFP behaved like the ΔCH1 and Δplakin constructs: it was enriched along MTs in distal axons accompanied by mild axon shortening and a trend towards increased spool formation (Figs.1G’’,K,L). In contrast, Shot-PE-Life::GFP localised strongly in GCs but also along axons (Figs. 1H’’, 3 and S4) and caused axon shortening and spool induction to similar degrees as Shot-PE::GFP (Fig.1K,L). However, other subcellular features were strikingly novel: (1) 38% of Shot-PE-Life::GFP-induced MT spools in GCs had a ‘tennis racket’ appearance with many MTs projecting diffusely through the centre of spools (Fig.3A and ‘white arrows’ in Fig.S4); (2) a number of neurons showed unusual MT bundles in close proximity to the cortex in the cell bodies (Fig.3D and ‘open curved arrows’ in Fig.S4); (3) about 60% of axonal MT bundles were split into two parallel portions that were decorated with strong Shot-PE-Life::GFP staining, and closely accompanied by F-actin staining that was unusually strong for axon shafts (Fig.3B,C and ‘white arrowheads’ in Fig.S4); these constellations suggested that the hybrid construct firmly cross-links and alters the sub-cellular arrangement of MTs and F-actin whilst taking on an unusual localisation itself (Figs.3 and S4; see Discussion). The aberrant localisation of Shot-PE-Life::GFP and its dominant MT phenotypes were clearly abolished when treating neurons with LatA, thus demonstrating the F-actin dependence even of this powerful hybrid construct (Fig.3E-G).

**Fig.3.**
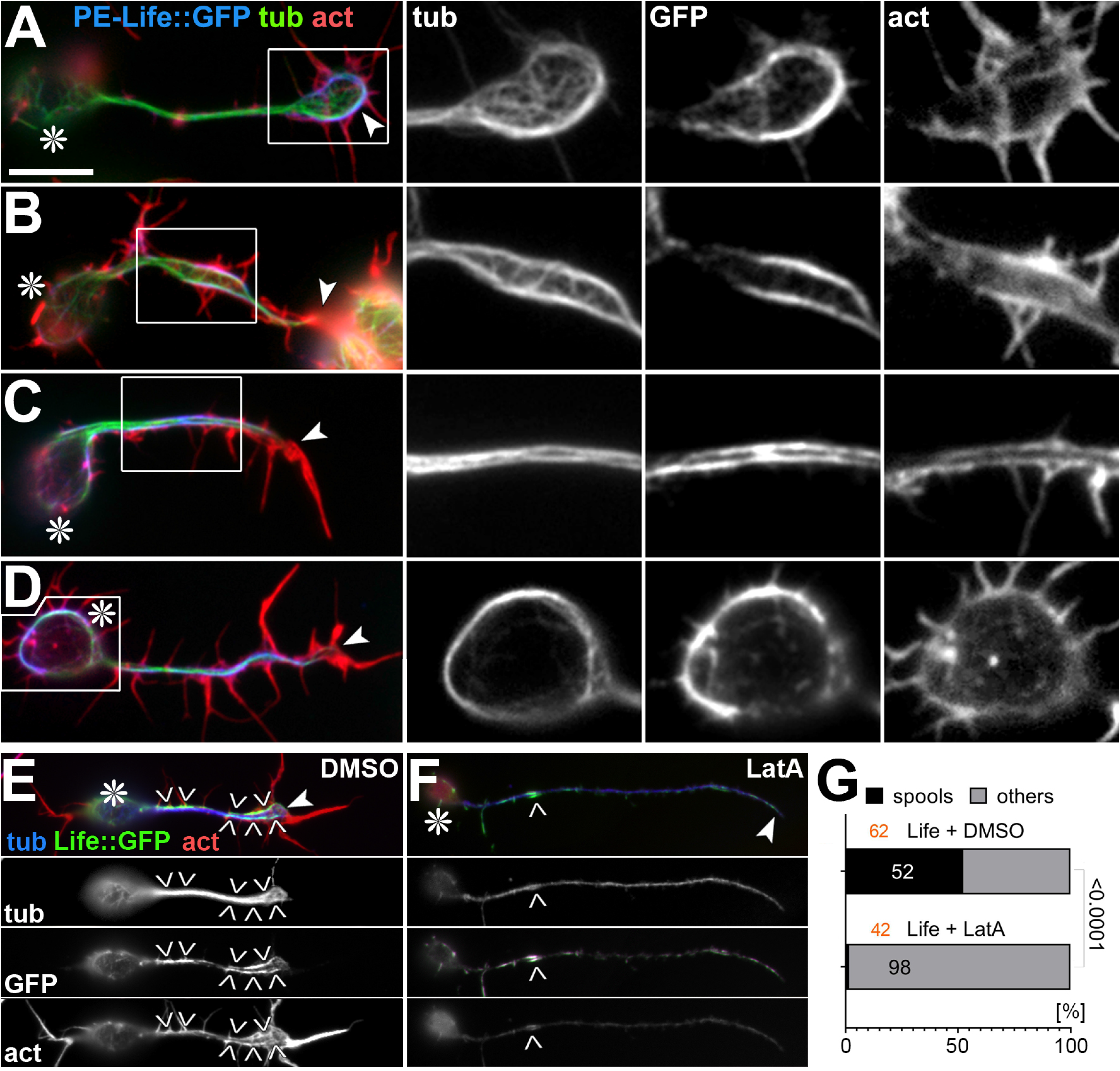
Characteristic phenotypes induced by Shot-PE-Life::GFP expression. **A-D**) Primary neurons at 6-8 HIV on glass with *scabrous-Gal4*-induced expression of Shot-PE-Life::GFP, stained for tubulin (green), actin (red) and GFP (blue); boxed areas are shown as twofold magnified single channel grayscale images on the right, as indicated. **E,F**) Shot-PE-Life::GFP-expressing neurons treated with vehicle (E) or latrunculin A (LatA; F), stained for the same markers as above but colour-coded differently (as indicated); grayscale images below show single channels. Asterisks in A-F indicate cell bodies, arrowheads tips of axons, chevrons in E and F indicate areas of high GFP concentration, and the scale bar in A represents 10 µm in all RGB images of A-D, 5 µm in grayscale images of A-D, and 20 µm in E. **G**) Percentage of Shot-PE-Life::GFP-expressing neurons showing spools (black) when treated with vehicle or LatA; number of analysed neurons in orange, percentage shown in bars, the X^2^ test result on the right.

Taken together, our GOF analyses suggest that the quality and quantity of F-actin networks can regulate Shot’s MT bundle-inducing function. The low affinity of Shot’s CH domains seems ideally tuned to read those differences in F-actin: high abundance of F-actin induces spools in GCs, and increases in Shot’s F-actin affinity (Shot-PE-Life) cause bundle modifications (split bundles) even in axon shafts (where F-actin networks are usually sparse; Qu et al., 2017; Xu et al., 2013).

### Shot’s axon length regulation involves MT spool formation in GCs and MT bundle maintenance

Our key readout for Shot GOF was the formation of MT spools in GCs. MT spools have been suggested to inhibit axon growth (Dent et al., 1999; Sánchez-Soriano et al., 2010). Accordingly, we find a strong negative correlation between spools and axon lengths when plotting the data from our over-expression experiments (black dots in Fig.4A); also neurons without Shot GOF plot onto this curve (Fig.4F,G and orange dots in A), including untreated wild-type neurons, neurons treated with LatA (less spools, enhanced axon length), or neurons lacking the F-actin-promoting factor Chickadee (Chic, the sole profilin in *Drosophila*; Gonçalves-Pimentel et al., 2011; slightly less spools, modest increase in axon length). Also spool formation in neurons without Shot GOF seems to be mediated by Shot, as suggested by *shot* mutant neurons where spool numbers are strongly reduced (Fig.4F; Sánchez-Soriano et al., 2010).

**Fig.4.**
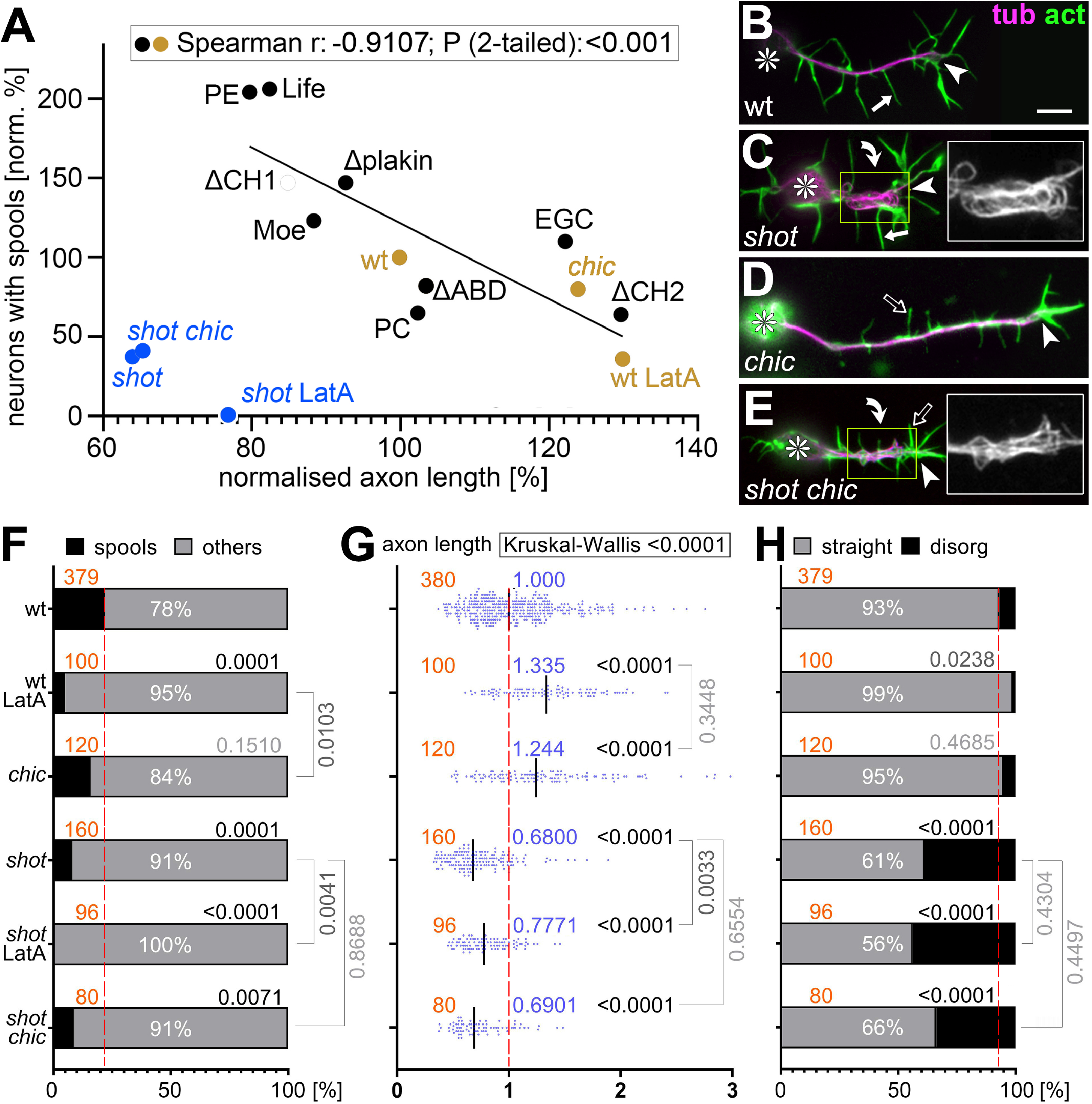
MT loops correlate with axon lengths but Shot has additional axon shaft phenotypes. **A**) Spearman correlation analysis comparing axon length and spool frequency. Black dots represent data from Fig.1K plotted against data from Fig.1L, and orange/blue dots match data from F and G; significant negative correlation (r- and P-values) for orange and black dots are shown in box at top. **B-E**) Primary neurons at 6-8 HIV on glass which are either wild-type (B), *shot^3/3^* (C), *chic^221/221^* (D) or *shot^3/3^ chic^221/221^* (E), stained for tubulin (magenta) and actin (green); asterisks indicate cell bodies, arrowheads tips of axons, curved arrows areas of MT curling, and white/open arrows normal/short filopodia (see quantifications in Fig.S5); yellow-boxed areas presented as twofold magnified insets showing the tubulin channel in grayscale; the scale bar in B represents 10 µm in all RGB images and 5 µm in insets. **F-H**) Quantification of neurons displaying MT spools in GCs (F), of axon lengths (G) and of neurons displaying MT curling in axonal shafts (H); numbers of analysed neurons are indicated in orange; median values in blue (G), percentages as white numbers within columns (F,H); P-values obtained via Mann–Whitney rank sum tests (G) or Chi^2^ tests (F,H) are shown in black/grey above bars or plotted data; all data were normalised to wild-type controls performed in parallel to all experiments (dashed red lines).

However, *shot* mutant neurons do not plot onto the correlation curve (blue dots in Fig.4A): instead of showing axon extension that would usually correlate with the absence of spools, their axons are very short. Furthermore, combinatorial studies revealed that the short axon phenotype of *shot* overrides LatA- or *chic*-induced axon elongation (Fig.4E,G). These short axon phenotypes of *shot* seem to mirror the occurrence of MT disorganisation in *shot* mutant neurons, where axonal bundles lose their parallel arrangements and take on curled, criss-crossing appearances (referred to as MT curling; Fig.4C). Like the axon length phenotype, axonal MT curling is not influenced by LatA treatment or loss of Chic (Fig.4E,G,H), thus demonstrating a further parallel between both phenotypes.

### Shot seems to work through two redundant mechanisms in MT bundle maintenance

Previous work has demonstrated that Shot prevents MT curling through a F-actin/Eb1/MT guidance mechanism: via its N-terminus it binds cortical F-actin and via its C-terminus to MTs and Eb1 - thus guiding the extension of polymerising MTs along the axonal cortex into parallel bundles; this F-actin/Eb1/MT guidance mechanism is supported by numerous structure-function, loss-of-function, pharmacological and genetic interaction studies (details in Fig.5; Alves-Silva et al., 2012; Hahn et al., 2021; Qu et al., 2019; Sánchez-Soriano et al., 2009).

**Fig.5.**
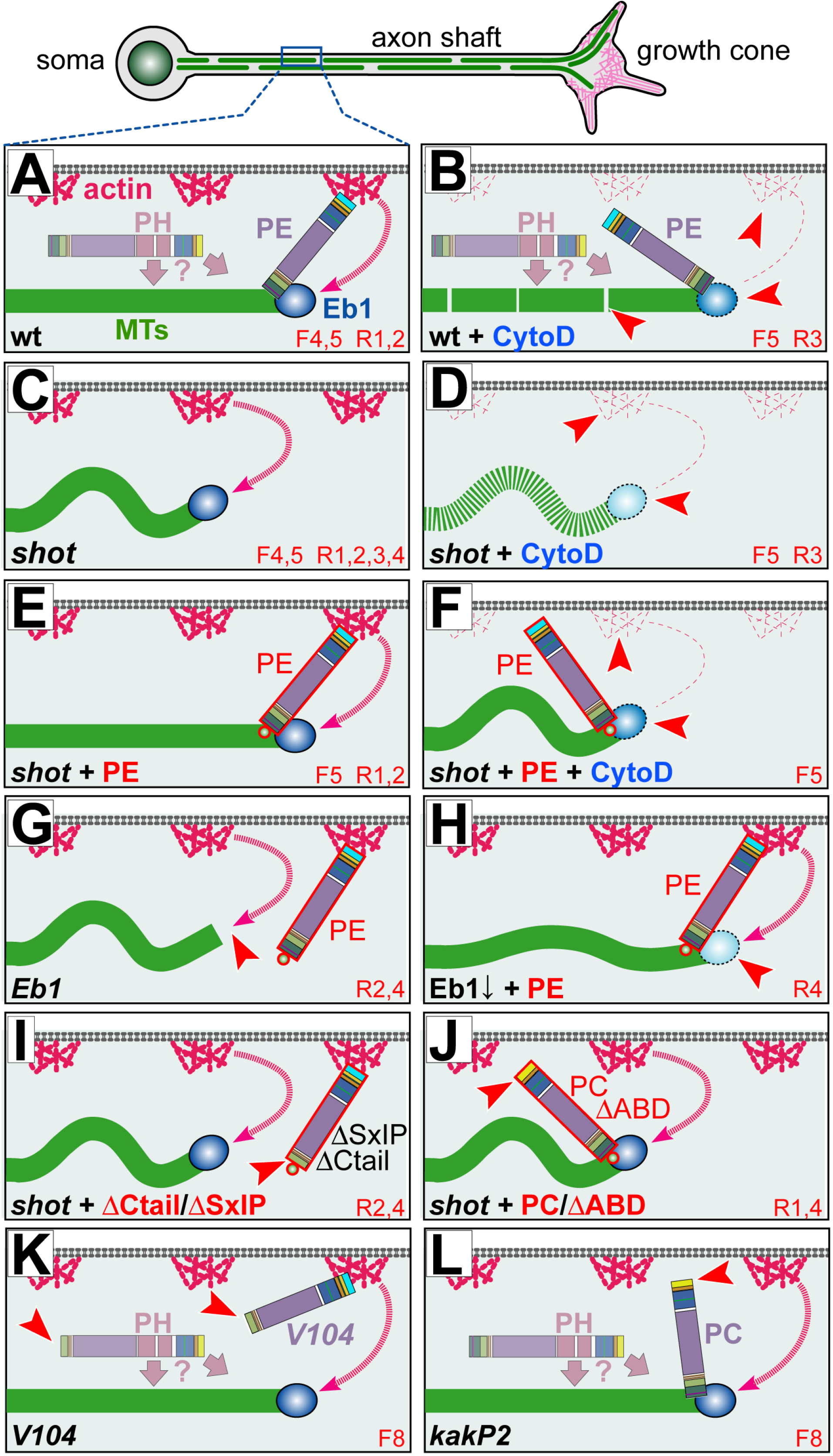
Schematic overview of existing experiments addressing Shot roles in MT bundle organisation. **A**) Schematic section of the axonal surface including cortical actin (magenta) anchoring the Shot N-terminus of CH1-containing isoforms (here PE) and promotes MT polymerisation (dashed magenta arrow); via its C-terminus, Shot-PE binds EB1 (dark blue) and MTs (green) thus cross-linking polymerising MT tips to the cortex and guiding their extension into parallel bundles; the PRR-containing PH isoforms (shown in pale) does not bind F-actin but we propose that it contributes to MT bundle formation/maintenance through yet unknown mechanisms (’?’; see Discussion). **B-L**) Different experimental conditions and their impact on MT behaviours; red numbers at bottom right indicate the information source: ‘F’ refers to figure numbers in this publication, ‘R’ indicates external references: (1) (Sánchez-Soriano et al., 2009), (2) (Alves-Silva et al., 2012), (3) (Qu et al., 2017), (4) (Hahn et al., 2021); red arrow heads point at specific lesions. Explanations: in wild-type neurons, CytoD eliminates cortical actin and weakens MT polymerisation (pale Eb1 with dashed outline), not strong enough to affect parallel MT arrangements but leading to MT gaps (B); in the absence of Shot, MTs curl (C) and MT networks shrink (they become vulnerable to lack of actin-promoting effects; D); guiding function is fully re-instated by targeted expression of Shot-PE (E; constructs red encircled with a green GFP dot at their ends); Shot-PE fails to guide MTs in the absence of actin, but it protects MT polymerisation (F); Eb1 deficiency eliminates MT guidance (G); MT curling upon reduced Eb1 levels (Eb1↓) can be rescued with Shot-PE expression (H); MT curling caused by loss of Shot (or Eb1; see Ref.4) cannot be rescued with Shot-PE variants that lack Ctail or Eb1-binding SxIP motifs (I; see Fig.7C) or the CH1 domain (J); absence of the same domains in *shot^V104^* (K) or *shot^kakP2^* (L) does not cause MT curling. We propose that the presence of the Shot-PH isoform (faintly shown in A,B,K,L) protects axons against loss of actin or F-actin/MT/Eb1 guidance mechanism, i.e. conditions which cause severe curling in the other experimental settings (C,F,G,I,J).

The F-actin/Eb1/MT guidance mechanisms would predict that removal of cortical F-actin from wild-type neurons (which can be achieved with the F-actin-inhibiting drug CytoD, but less so with LatA or loss of Chic; Qu et al., 2017) should mimic the *shot* mutant MT curling phenotype. However, CytoD application to wild-type neurons fails to cause MT curling; instead it causes a deficit in MT polymerisation leading to gaps in MT bundles (Figs.S1B, 6B, 5B; Qu et al., 2017) – which may also explain why loop suppression upon CytoD staining (Fig.S1) does not enhance axon growth as observed with LatA (Sánchez-Soriano et al., 2010).

The fact that CytoD fails to mimic the MT curling phenotype observed in *shot* mutant neurons (Fig.6B *vs.* C) might indicate that F-actin/Eb1/MT guidance is not the only mechanism through which Shot contributes to MT bundle maintenance. For example, Shot might work through further isoforms beyond Shot-PE (the only isoform shown so far mediating guidance; Fig.5E,F,H-J). To test this possibility, we used Shot-deficient mutant neurons in which the MT curling phenotype was rescued by the F-actin/Eb1/MT guidance mechanisms, i.e. the expression of Shot-PE (Figs.5E, 6E). When these seemingly normal neurons were treated with CytoD, strong MT curling was induced (Figs.6F, 5F), suggesting that these neurons lack some actin-independent bundle-maintaining functions of Shot that are present in wild-type neurons.

**Fig.6.**
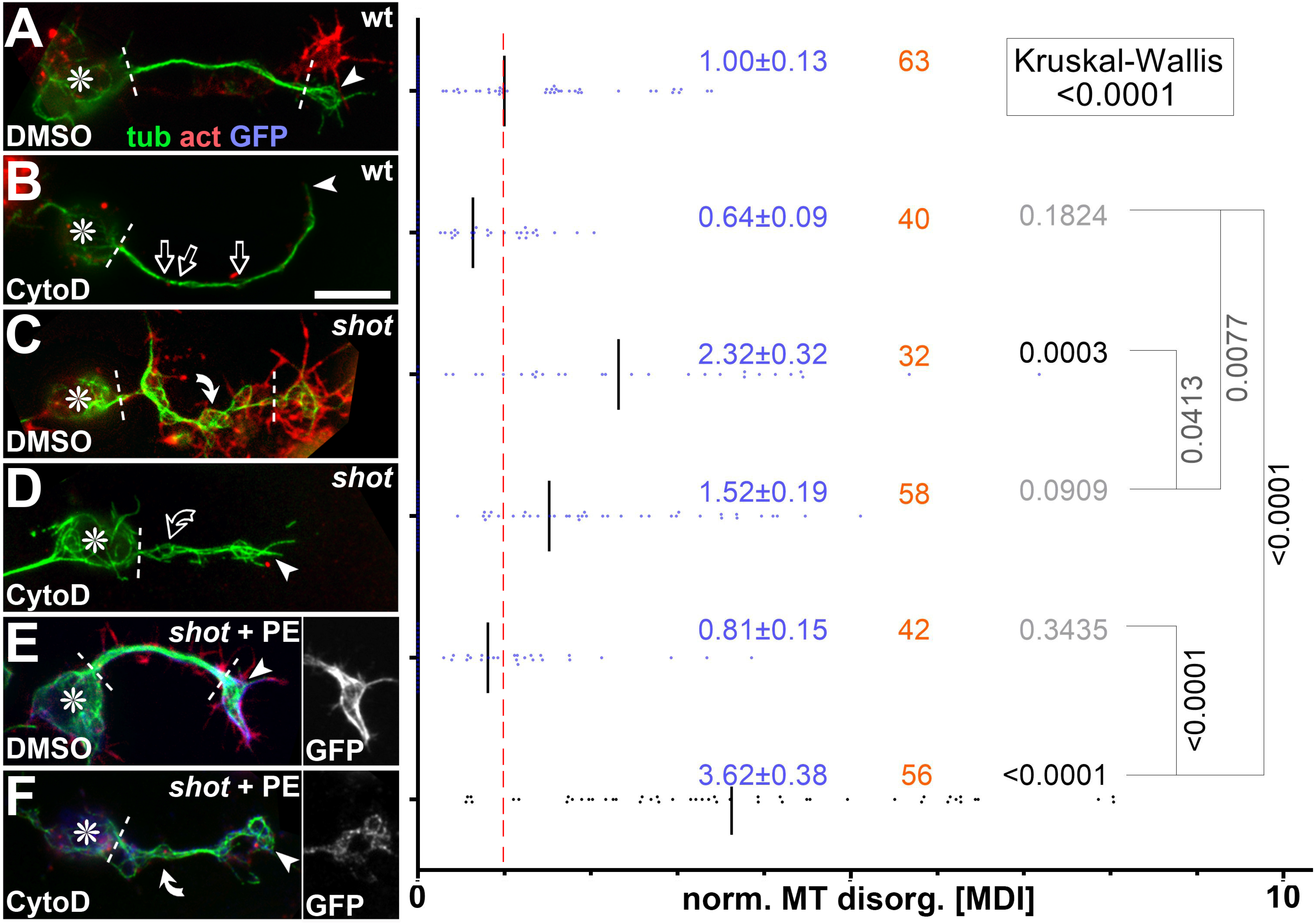
CytoD experiments confirming the F-actin-dependent guidance mechanism of Shot. **Left side**: Primary neurons of different genotypes (as indicated: wt, wild-type; *shot*, *shot^3/3^*; *shot* + PE, *shot^3/3^* expressing Shot-PE) at 6-8 HIV on ConA, treated with vehicle (DMSO) or cytochalasin D (CytoD) as indicated, and stained for tubulin (green), actin (red) or GFP (blue); asterisks indicate cell bodies, arrowheads the tip of axons, white lines demarcate the axon shaft, open arrows gaps in axonal tubulin bundles and white/open curved arrows areas of normal/fractured MT curling; scale bar in B represents 10 µm in all images. **Right side:** Quantification of the degree of MT curling in the axon shafts (between white dashed lines or dashed line and arrow head in images on the left) of each genotype, measured in MDI and normalised to wildtype controls (red dashed line); numbers of neurons analysed are indicated in orange, mean ± SEM in blue and results of Mann-Whitney rank sum tests are shown in black/grey. Further explanations are given in Fig.5.

F-actin/Eb1/MT cross-linkage requires the CH1 and Ctail domains of Shot, as revealed by rescue experiments in *shot* mutant neurons (using Shot-PC and Shot-PE-ΔCtail; Fig.5I,J). These two domains are specifically missing in *shot^kakP2^* and *shot^V104^* mutant alleles, which do not affect the rest of the endogenous *shot* gene locus (details in Figs.1A and 7; Bottenberg et al., 2009; Gregory and Brown, 1998). The two alleles should therefore eliminate the guidance mechanism, but might retain the other bundle-maintaining function of Shot (’PH’ in Figs.4 I,J *vs.* K,L).

When analysed in whole embryos, both mutant alleles clearly caused partial loss-of-function mutant phenotypes: *shot^kakP2^* strongly affected the nervous system (Bottenberg et al., 2009; Gregory and Brown, 1998), and *shot^V104^* defects seemed to restrict to non-neuronal tissues (Fig.S6). When cultured as primary neurons, we measured the degree of MT curling in the axon shaft, which is the area where the guidance mechanism is expected to make its prime contributions. For *shot^V104^* we found no obvious phenotype, *shot^kakP2^* revealed only a trend, whereas the *shot^3^* null mutant alleles displayed severe MT curling along axon shafts (Fig.8A-D,F). For *shot^V104^* mutant neurons we repeated the experiment culturing them on concanavalin A which is a more challenging condition causing greater mechanical strain (Prokop et al., 2012). When challenged this way, *shot^V104^* mutant neurons displayed robust MT curling. This suggests that loss of the F-actin/MT/Eb1 guidance mechanism weakens the overall machinery of MT bundle maintenance: under modest conditions, its absence can be masked by the other functions of Shot, but not when mechanically challenged.

**Fig.7.**
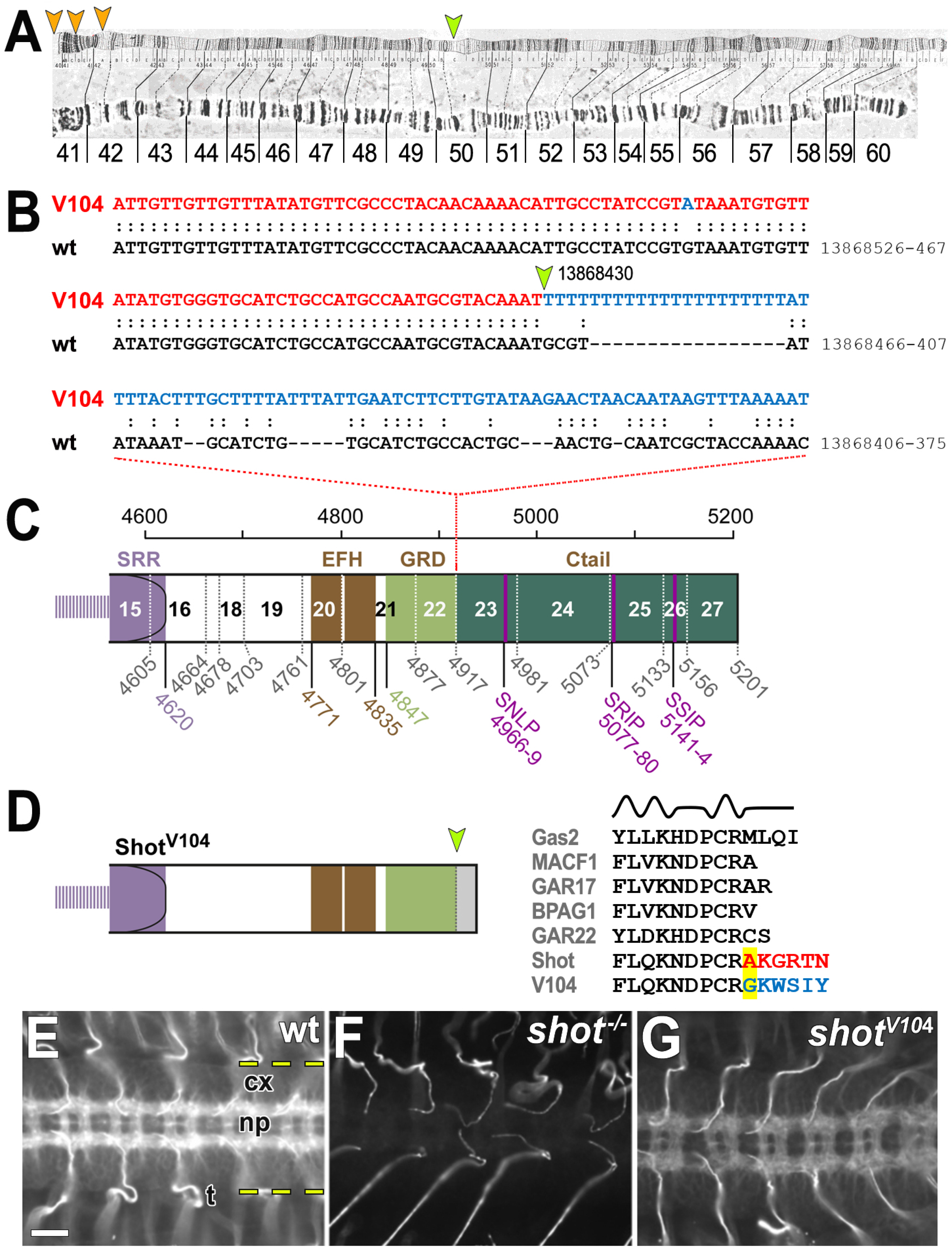
The *shot^V104^* breakpoint removes the Ctail. **A**) View of the 2R polytene chromosome (Lindsley and Zimm, 1992) indicating the mapped breakpoint in 50C (orange arrow) and potential sites of the second breakpoint in the centromeric region of 2R (orange arrowheads) suggested by the mapping positions of several clones with matching sequences (when using the BLAST function in flybase.org and the blue sequence in B as query); clones with matching sequences: *DS03708* (42A4-42A5), *BACR04E10* (41C-41D), *BACR07J16* (41C-41C), *BACR05A24* (41C-41D), *BACR05A24* (41C-41D), *BACR03D04* (40D-40D). **B**) Alignment of the wild-type and *V104* mutant genomic sequences of *shot* indicating the breakpoint (yellow arrow) in position 13,868,412 (primary assembly 2R: 13,864,237-13,925,503 reverse strand) and the newly fused sequence in *shot^V104^* (blue) likely derived from the other end of the inversion that would usually be situated near the position of the second breakpoint (orange arrow in A). **C**) Schematic of the Shot-PE protein (FBtr0087618) drawn to scale and indicating domain/motif borders (coloured numbers below; compare Fig.1A) as well as exon borders (stippled vertical lines, grey numbers, exon numbers indicated between lines); *V104* the breakpoint is situated in intron 22/23. **D**) The predicted V104 protein is truncated behind the GRD (yellow arrow) potentially reading into intronic sequences (grey). Comparison of the V104 sequence at the breakpoint (highlighted yellow) with sequences of GRDs from normal Shot and other GRD-containing proteins (listed in grey; taken from Alves-Silva et al., 2012) strongly suggest that the truncation does not affect the final α-helix and amino acid changes occur behind the GRD. **E-G**) Ventral nerve cords of stage 16 embryos (cx, cortex containing cell bodies; np, neuropile containing synapses and as -/descending tracts; both separated by dashed yellow lines) stained with the Shot-C antibody against the C-terminal part of the spectrin repeat rod (Fig.1A; Strumpf and Volk, 1998); staining reveals the presence of protein in wild-type (E), absence in homozygous *shot* null mutant embryos (F) and presence in hemizygous *shot^V104/MK1^* mutant embryos where reduced expression is due to the absence of one gene copy (*V104* is over the *MK1* deficiency); scale bar in E represents 20 µm in E-G.

**Fig.8.**
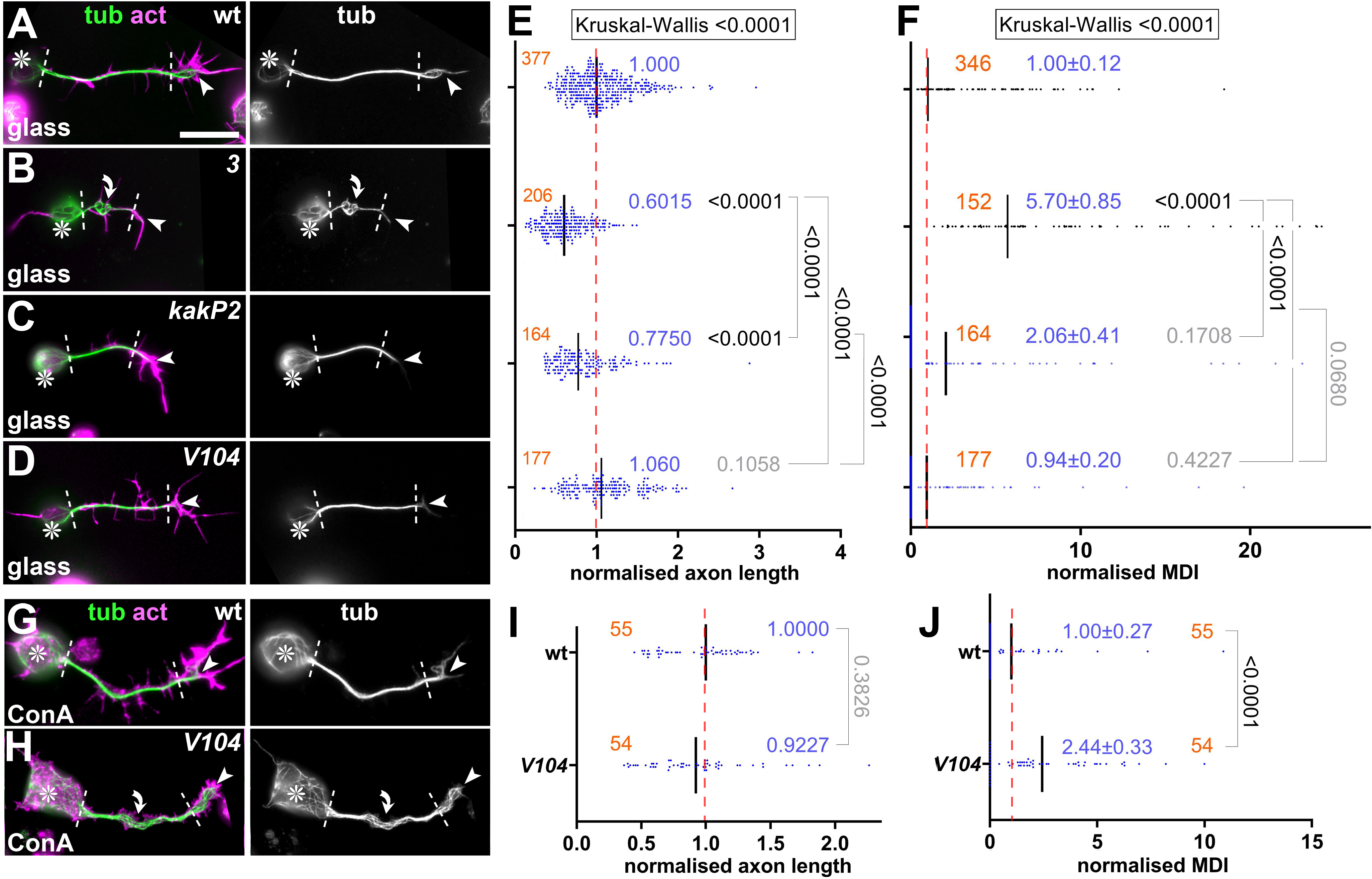
Phenotypes of *shot^kakP2^* and *shot^V104^* mutant primary neurons. **A-D,G,H**) Images of neurons at 6-8 HIV of different genotypes (wt, wild-type; *3*, *shot^3/3^*; *kakP2*, *shot^kakP2/kakP2^*; *V104*, *shot^V104/Df(MK1)^*) cultured on glass (A-D) or ConA (G,H) and stained for tubulin (green), actin (red) or GFP (blue); greyscale images on the right show only the tubulin channel; asterisks indicate cell bodies, arrowheads the tips of axons, white lines demarcate axon shafts, curved arrows areas of MT curling; scale bar in A represents 20 µm in A-D and 10 µm in G,H. **E,F,I,J**) Quantifications of axon length (E,I) and MT curling (measured in MDI; F,J), both normalised to wild-type controls (red dashed line); numbers of neurons analysed are indicated in orange, mean ± SEM in blue and results of Mann-Whitney rank sum tests are shown in black/grey.

## Discussion

### Neuronal roles of Shot involve isoform-specific actin-dependent and -independent functions

Spectraplakins are well conserved across the animal kingdom; they are essential cytoskeletal regulators in neurons, linked to severe MT curling in mammals and *Drosophila* alike (Voelzmann et al., 2017). Many mechanistic insights were gained using *Drosophila* Shot as a model, and F-actin/MT linkage has emerged as a central theme that is consistent also with roles in non-neuronal cells (Kodama et al., 2003). Here we refined our understanding of Shot’s actin dependency during MT regulation, whilst also proposing the co-existence of actin-independent functions involved in MT bundle promotion.

### Shot’s roles in spool formation are regulated by F-actin

Our findings suggest that F-actin is an important instructor of Shot’s MT-regulating roles. For example in GCs, Shot is an essential regulator of spool formation in an F-actin-dependent manner: (1) it can be suppressed when depleting F-actin (LatA, CytoD; Sánchez-Soriano et al., 2010), (2) when changing the properties of F-actin networks (CK666), or (3) when changing Shot’s actin-binding properties as observed with Shot-PC, Shot-PE-ΔABD, Shot-PE-ΔCH2, Shot-PE-Moe and Shot-PE-Life. In contrast, in the axon shaft, F-actin networks are far less prominent (Xu et al., 2013), which seems sufficient to support cortical guidance of MT polymerisation but not enough to induce prominent changes to MT bundles even when overexpressing Shot-PE. In contrast, Shot-PE-Life was able to induce abnormal MT bundle split in the shaft, suggesting that increased F-actin affinity is sufficient to tip the balance in an F-actin-sparse environment and change the MT-regulating behaviour of Shot.

Taken together, these experiments suggest that proper Shot function requires well-balanced interaction with F-actin networks, and the spectacular phenotypes we observe with Shot-PE-Life (Figs.3 and S4) suggest, that our findings can be turned into new genetic tools to investigate how changes in the cytoskeleton impact on neuronal architecture, dynamics and even physiology.

Our experiments with Shot-PE-Life have demonstrated a clear F-actin-dependence of the induced MT phenotypes (Fig.3F,G). They also suggested that this construct was able to induce ectopic F-actin in axon shafts (Figs.3, S4), potentially reflecting mutual regulation mediated through Shot. This may involve known roles of the Shot C-terminus in promoting F-actin nucleation (Sánchez-Soriano et al., 2009), thus creating a scenario in which the strong localisation of Shot-PE-Life along axon shafts might trigger a positive feedback loop by nucleating more F-actin which then enhances Shot-PE-Life localisation.

In normal Shot-PE, direct binding through the CH domains might not be sufficient to trigger changes in Shot function, and also the plakin domain appears functionally involved. To our knowledge, the only plakin domain-binding factors reported so far are transmembrane adhesion factors including integrins and collagen XVII at mammalian hemidesmosomes (Aumailley et al., 2006) and potentially the N-CAM homologue Fasciclin II in *Drosophila* neurons (Voelzmann et al., 2017). Since the localisation of such adhesion factors is dependent on F-actin (Woichansky et al., 2016), they might provide a potential second route through which F-actin can influence Shot activity.

In summary, we have built a case for regulatory impacts of F-actin networks on Shot function which, in turn, trigger MT network changes that impact on axon growth; this is best exemplified by the negative correlation between spool formation and axon growth (Fig.4A; Dent et al., 1999; Sánchez-Soriano et al., 2010).

### Shot displays prominent F-actin-independent roles in axons

Shot also plays major roles in maintaining MT bundles in axon shafts. We confirmed here the importance of F-actin/MT/Eb1 cross-linkage for MT guidance into parallel bundles (Alves-Silva et al., 2012; Hahn et al., 2021; Figs.6, 8). We believe that these roles are merely permissive and not subject to F-actin dependent regulation, because F-actin networks in axon shafts appear sparse and far less dynamic when compared to GCs. The key impact of MTs not staying in proper bundles is likely due to the fact that they cannot contribute with the same rigour to the growth events at GCs.

In addition to the guidance mechanism involving F-actin/MT/Eb1 cross-linkage, we also presented strong arguments for additional functions of Shot in MT bundle maintenance that are independent of this form of cross-linkage. Considering the enormous importance that MT bundles have for the long-term survival of axons, it appears only logical to have redundant mechanisms to maintain these bundles and prevent axonopathies (Prokop, 2021).

In our view, the best candidate to mediate such F-actin-independent functions of Shot is the unique Shot-PH isoform. Shot-PH is highly expressed in the nervous system, has a C*-type N-terminus (non-F-actin-binding like Shot-PC; Fig.1A), and stands out as the only isoform containing a large central PRR (plakin repeat region; Fig.S7; flybase.org reference: FBgn0013733; Hahn et al., 2016; Röper and Brown, 2003; Voelzmann et al., 2017).

PRRs are conserved in mammalian dystonin and ACF7/MACF1 (Voelzmann et al., 2017), but very little is known about their role or potential binding partners. PRRs of *Drosophila* Shot play regulatory roles at epithelial adherens junctions through unknown mechanisms (Röper and Brown, 2003). In mammals, the PRR-containing isoform MACF1b was shown to associate with the Golgi (Lin et al., 2005). However, it is difficult to imagine how Golgi-related mechanisms could maintain MT bundles in the absence of F-actin-dependent guidance mechanisms of Shot. In our view, investigating the potential roles and mechanisms of PRRs in axons would therefore have great potential to deliver new mechanisms that can advance our understanding of axon maintenance and architecture (Prokop, 2020).

As a first step to study PRRs, we generated flies carrying a CRISPR/Cas9-mediated PRR deletion. Unfortunately, *shot^ΔPRR^* mutant flies displayed unexpected splicing defects resulting in a strong loss-of-function mutant allele (details in Fig.S7); whilst being potentially interesting for molecular geneticists that work on splicing mechanisms, this allele was unsuitable for our purposes. An alternative strategy could be to identify PRR-binding or -associating proteins (Lin et al., 2021), and then use versatile *Drosophila* genetics in combination with our culture model (Prokop et al., 2013) to establish their potential involvement in bundle maintenance. Amongst the PRR-interacting proteins, we would expect to find also Eb1-binding proteins or even Eb1 itself (note that PRR contains a potentially Eb1-interacting SNLP motif as similarly found in the Ctail; Fig.7C; Honnappa et al., 2009); a link from the PRR to Eb1 could explain an important conundrum posed by the current data: loss of Eb1 causes MT curling, but the deletion of the Eb1-binding Ctail from all Shot isoforms does not (Fig.5G vs. K) – the PRR might be the missing puzzle piece.

Taken together, we propose a system of redundant Shot-mediated mechanisms that promote axonal MT bundle architecture - in addition to other factors expected to be involved, such as classical MAPs or mitotic kinesins (Guha et al., 2021; Hahn et al., 2019; Prokop, 2020). Such robust redundancy makes sense when considering the enormous importance of these MT bundles for axonal longevity (Prokop, 2021). We believe that the study of Shot-PH can establish new investigative paths towards a more profound understanding of axon architecture, thus bridging a gap in the field that may provide important explanations for a wide range of axonopathies and new avenues for their treatment.

## Materials and Methods

### Fly strains

The following fly stocks were used: Oregon R as wild-type control and the strong loss-of-function or null alleles *chic^221^* (Verheyen and Cooley, 1994)*, shot^3^* (Kolodziej et al., 1995), *shot^kakP2^* (synonymous to *P{lacW}shot^k03405^*; Gregory and Brown, 1998), *shot^HG25^* (Prokop et al., 1998) and *shot^V104^* (Strumpf and Volk, 1998). All mutant stocks were kept and selected with *twi-Gal4/UAS-GFP* green balancers (Halfon et al., 2002). Existing transgenic lines we used included the *scabrous-Gal4*, *eve-Gal4^RN2E^* and *stripe-Gal4* driver lines (Fujioka et al., 1999; Mlodzik et al., 1990; Subramanian et al., 2003), *UAS-mCD8::GFP* (Luo et al., 1994), *UAS-shot-RE-GFP and UAS-shot-RC-GFP* (Lee and Kolodziej, 2002), *UAS-EGC-GFP* (Subramanian et al., 2003), *UAS-shot-RE-Δplakin-GFP* (Bottenberg et al., 2009) and *UAS-Act5C-GFP* (Bloomington Stock Center; Kelso et al., 2002).

### *Drosophila* primary neuronal cell culture

Neuronal cell cultures were generated as detailed elsewhere (Prokop et al., 2012; Voelzmann and Sánchez-Soriano, 2021). Embryos were dechorionated for 1.5 min in 50% domestic bleach, correct stages (usually stage 11; Campos-Ortega and Hartenstein, 1997) and genotypes were selected under a fluorescent dissecting microscope, transferred to sterilised centrifuge tubes containing 100µl of 70% ethanol, washed in sterile Schneider’s medium containing 20% fetal calf serum (Schneider’s/FCS; Gibco) and, eventually, homogenised with micro-pestles in 1.5 ml centrifuge tubes containing 21 embryos per 100 μl dispersion medium (Prokop et al., 2012). They were left to incubate for 4 min at 37°C. Dispersion was stopped with 200 μl Schneider’s/FCS, cells were spun down for 4 mins at 650 g, supernatant was removed and cells were re-suspended in 90 µl of Schneider’s/FCS; 30 μl drops were placed in culture chambers and covered with cover slips. Cells were allowed to adhere to cover slips for 90-120 min either directly on glass or on cover slips coated with a 5 µg/ml solution of concanavalin A, and then grown as a hanging drop culture at 26°C usually for 6-8 hrs.

Transfection of *Drosophila* primary neurons was executed as described previously (Qu et al., 2019). In brief, 70-75 embryos per 100 μl dispersion medium were used. After the washing step and centrifugation, cells were re-suspended in 100 μl transfection medium [final media containing 0.1-0.5 μg DNA and 2 μl Lipofectamine 2000 (L2000, Invitrogen)], incubated following manufacturer’s protocols (Thermo Fisher, Invitrogen) and kept for 24 hrs at 26°C. Cells were then treated again with dispersion medium, re-suspended in culture medium and plated out as described above.

### Drug application and immunohistochemistry

For drug treatments, solutions were prepared in cell culture medium from stock solutions in DMSO. Cells were treated for 4 hrs with 200 nM latrunculin A (Biomol International), 0.4 μg/ml cytochalasin D (Sigma) or 100 nM CK666 (Sigma), respectively. For controls, equivalent concentrations of DMSO were diluted in Schneider’s medium.

Culture medium was carefully removed and cells fixed for 30 mins with 4% paraformaldehyde in 0.05 M phosphate buffer (pH 7-7.2), then washed in PBT (phosphate buffered saline with 0.3% TritonX-100). Incubation with antibodies was performed in PBT without blocking reagents. The following antibodies were used: anti-α-tubulin (clone DM 1A, 1:1000, mouse, Sigma), anti-Shot raised against aa3450-4714 (C-terminal end of the spectrin repeat region; guinea pig; 1:200; Strumpf and Volk, 1998); anti-GFP (1:500, goat, Abcam), and FITC-, Cy3 - or Cy5-conjugated secondary antibodies (1:200, purified from donkey, Jackson Immunoresearch). F-actin was stained with TRITC- or Cy5-conjugated Phalloidin (Sigma; 1:100). Coverslips with stained neurons were mounted on slides using Vectashield medium (Vector labs) or ProLong Gold Antifade Mountant (ThermoFisher Scientific).

### Stage 17 embryo dissections

Dissection of late stage 17 embryos (stages according to Campos-Ortega and Hartenstein, 1997) was carried as described in great detail elsewhere (Budnik et al., 2006). In brief, embryos were dissected flat in PBS on Sylgard-coated cover slips with the help of sharpened tungsten needles and Histoacryl glue (Braun, Melsungen, Germany), followed by 1 hr fixation in 4% paraformaldehyde, 1 hr wash in PBT and the same histochemical staining steps as mentioned above using the following antibodies: anti-FasII (1D4 2F3, DSHB; mouse, 1:20; Van Vactor et al., 1993), anti-GFP (see above) and anti-Synaptotagmin (rabbit polyclonal; 1:1,000; Littleton et al., 1993). Embryos were cut out from the glue using razor blade splinters or the tungsten needles and embedded in glycerol.

### Imaging and image analysis

Standard imaging was performed with AxioCam 506 monochrome (Carl Zeiss Ltd.) or MatrixVision mvBlueFox3-M2 2124G digital cameras mounted on BX50WI or BX51 Olympus compound fluorescent microscopes. Measurements from images were carried out using ImageJ (segmented line and freehand selection tools). Only neurites at least twice the length of the soma diameter were analysed using α-tubulin staining and measuring from the edge of the cell body to the tips of the axons (excluding MTs in filopodia); in cases where neurites branched the longer branch was measured, in cases where 2 neurites extended from a single cell the longer value was taken. The degree of disorganised MT curling in axon shafts was established either as binary readout (% of neurons with disorganisation) or as “MT disorganisation index” (MDI) described previously (Qu et al., 2019; Qu et al., 2017); in short: the area of disorganised curling was measured with the freehand selection in ImageJ; this value was then divided by axon length (see above) multiplied by 0.5 μm (typical axon diameter, thus approximating the expected area of the axon if it were properly bundled); in this study, MDI measurements were restricted to the axon shaft, i.e. from the cell body to the base of GCs (white dashed lines in Figs. 6, 8). Filopodia numbers were counted per neurite. GCs containing looped MT bundles (spools) were classified according to previous publications (Sánchez-Soriano et al., 2010). Graphpad Prism was used to describe data and perform statistical tests. Data were usually not normally distributed, and the median was determined for axon length; since MDI measurements contain many zero-value data, the mean and standard error of the mean (SEM) had to be used to obtain meaningful numbers. For statistical analyses, the Chi-square test was used when comparing percentages, Kruskal–Wallis one-way ANOVA test to compare groups, and Mann– Whitney Rank Sum Tests (indicated as P_MW_) to compare pairs of data. For the correlation, r and p-value were determined via non-parametric Spearman correlation analysis (tests showed that data are not distributed normally).

### Electron microscopy

Procedures followed protocols published in detail elsewhere (Budnik et al., 2006). In brief, embryos were injected with 5% glutaraldehyde in 0.05 M phosphate buffer, pH 7.2, the injected specimens were cut open at their tips with a razor blade splinter, postfixed for 30-60 min in 2.5% glutaraldehyde in 0.05 M phosphate buffer, briefly washed in 0.05 M phosphate buffer, fixed for 1 h in aqueous 1% osmium solution, briefly washed in dH2O, treated en bloc with an aqueous 2% solution of uranyl acetate for 30 min, dehydrated, and then transferred to araldite or TAAB LV (TAAB Laboratories Equipment, Berkshire, UK). Serial sections of 30-50 nm (silver-grey) thickness were transferred to formvar-covered carbon-coated slot grids, poststained with lead citrate for 5-10 min, and then examined on a JEOL 200CX (Peabody, MA) or Hitachi H600 (Tokyo, Japan).

### Cloning of *shot* constructs

The CH deletions (ΔCH1, ΔCH2, ΔABD; *UAS-shot.RE-DeltaABD.GFP* now available at Bloomington, #93282) were made by PCR amplification of 2 DNA fragments flanking the CH domains, using respective primers listed in the table which contained homologous sequences to anneal them into a template for further PCR amplification. The PCR product was digested and ligated into *pET20b* vector (Novagen) using AscI and XhoI. To insert alternative actin-binding domains (Lifeact source: *pCMVLifeAct-TagGFP2* vector, Ibidi; Moesin was a gift from Tom Millard; Millard and Martin, 2008; *UAS-shot.RE-Lifeact.GFP* now available at Bloomington, #93283), they were amplified in parallel to the 2 CH domain-flanking sequences and annealed in triplet constellation for making the template. PCR amplification was used to add NotI/XbaI restriction sites to the 5’ and 3’ ends followed by digestion and ligation into a modified version of the *pUASp* vector (Invitrogen; kindly provided by Tom Millard) which confers ampicillin resistance and tags the construct N-terminally with eGFP (referred to as *pUASp-eGFP*). N-terminal constructs in *pUASp-eGFP* were amplified in chemically competent TOP10 *E. coli*. and used for transfection into primary neurons (see above).

For making the respective full-length Shot-PE constructs carrying the N-terminal variations (Shot-PE-ΔABD, Shot-PE-ΔCH1, Shot-PE-ΔCH2, Shot-PE-Life, Shot-PE-Moe), *Nterm_Recomb* primers were used to amplify the N-terminal constructs from the *pET20b* vector. These were then used to replace the *GalK* cassette in full-length shot-RE within *M-6-attB-UAS-1-3-4* vector via recombineering strategies (Alves-Silva et al., 2012) and the positive/negative selection strategy (Warming et al., 2005). The *GalK* cassette was originally inserted into *M-6-attB-UAS-1-3-4 shot-RE*-borne *shot-RE* by using similar recombineering steps with *GalK* which had been amplified with primers that added the same homology arms as mentioned above.

The completed constructs in *M-6-attB-UAS-1-3-4* vector were amplified in Epi300 competent cells (EpiCentre) in LB-Chloramphenicol medium, adding CopyControl solution (EpiCentre) 2 hrs before the miniprep. Amplified constructs were used to generate transgenic flies (outsourced to BestGene, Chino Hills, CA 91709, US) using PhiC31-mediated site-specific insertion using a specific attB landing site on the third chromosome (*PBac{y*^+^*-attP-3B}CG13800^VK00031^*; Bloomington line #9748; Alves-Silva et al., 2012). This same landing site was used for all constructs to avoid position effects and achieve equal expression levels of all constructs (Bischof et al., 2007).

**Tab. 1.**
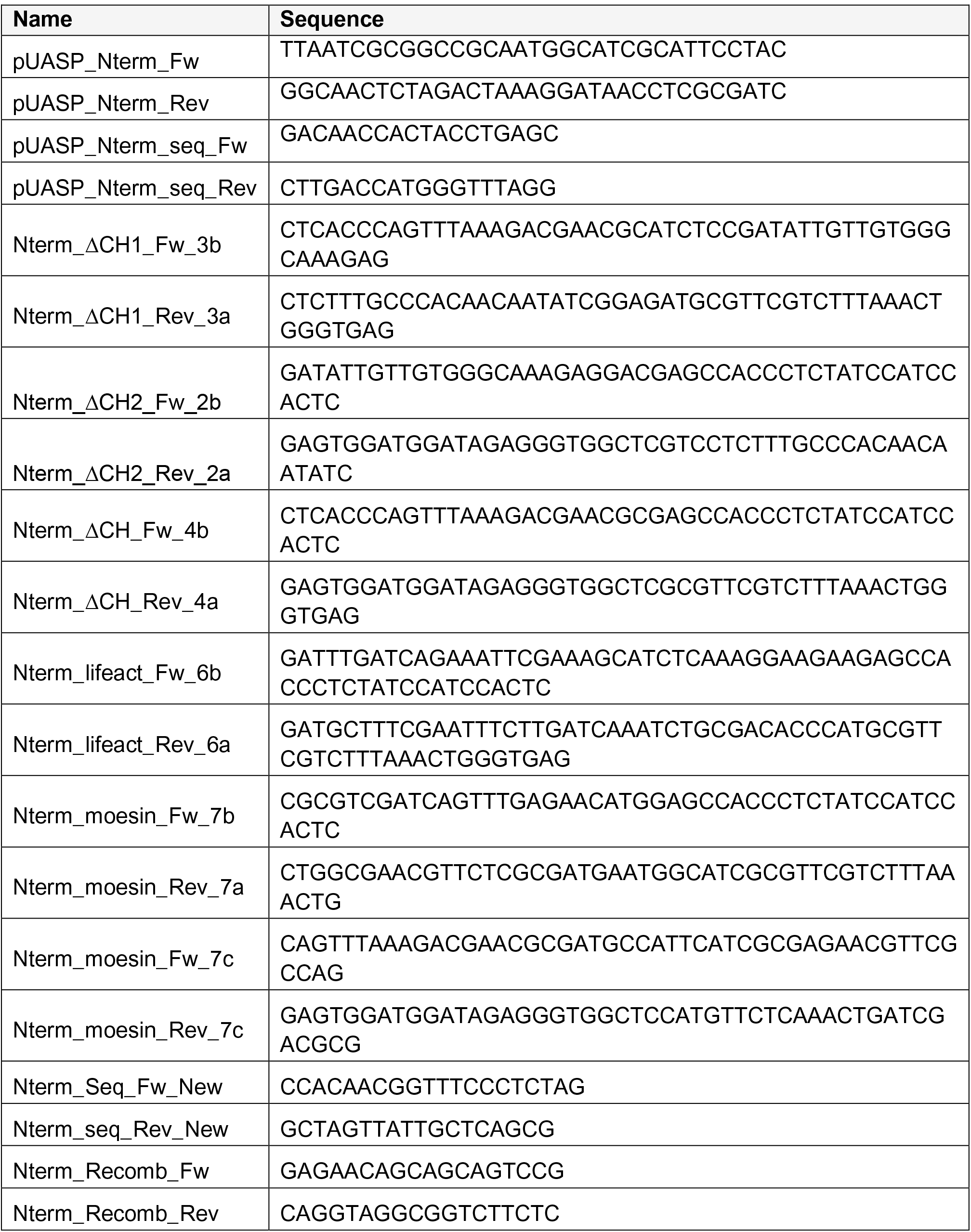
List of primers

### Generating *shot^ΔPRR^* mutant flies

The PRR domain (exon 12 of shot-RH, FBtr0087621) was excised from the *shot* genomic region and replaced with 3xP3-DsRed (driving DsRed expression in the eye) via CRISPR/Cas9 mediated homology-directed repair. Suitable gRNA target sites (5’ gRNA: GAGTGCTAACCTCCTGACTAG, 3’ gRNA: CTGTTCTGCCGGCAGGAGCAC) were identified by CRISPR optimal target finder (Gratz et al., 2014) and cloned into pCFD4-U6:1_U6:3tandemgRNAs (gift from Simon Bullock; Addgene plasmid # 49411; RRID:Addgene 49411) via Gibson assembly (NEB). Adjacent 2kb 5’ and 3’ homology regions were cloned into *pHD-DsRed-attP* (gift from Melissa Harrison & Kate O’Connor-Giles & Jill Wildonger, Addgene plasmid # 51019, RRID:Addgene_51019) 5’ region via EcoRI/NotI, 3’ region via BglII/PstI) using the following primer pairs:

- 5’ HR fwEcoRI: AAAAGAATTCctcgtttgttcgctcttaccc
- 5’ HR revNotI: AAAAGCGGCCGCCTGAAAGGATTCGATTAGAACTTTATTAG
- 3’ HR fwBglII AAAAAGATCTGTAAGTCTCAGAACACTCGAGG
- 3’ HR revPstI AAAACTGCAGTCGATCTCATCCTTGATTTGCTATTTAAAC

Constructs were injected into *M{Act5C-Cas9.P.RFP-}ZH-2A DNAlig4^169^* flies (Bloomington stock #58492) and selected for dsRed positive flies. Positive candidates were confirmed by sequencing.

### qRT-PCR analysis of *shot^ΔPRR^* mutant embryos

For RNA isolation, at least ten *Drosophila* third instar larvae were placed in Trizol (Invitrogen) and homogenised using a pestle. Total RNA was isolated using the NucleoSpin RNA II kit (Macherey & Nagel) and RNA concentration was analysed via a NanoDrop spectrophotometer (Thermo Scientific). For first strand cDNA synthesis, 500 ng of total RNA was transcribed using the QuantiTect RT Kit (Qiagen). Real-time PCR was performed with 1 µl cDNA per reaction using the Power SYBR Green PCR Master Mix (ThermoFischer Scientific) as detection dye. Experiments were performed with the BioRad C1000 Thermal Cycler. cDNA samples were run in triplicates, the average CT was used to analyse the expression levels via the −2ΔΔCT method. Experiments were repeated with independently isolated RNA samples. Actin 5C (Act5C, act) and Ribosomal protein L32 (RpL32, rp49) were used as reference genes. Expression analysis was performed using BioRad C1000 System software and GraphpadPrism. The following oligonucleotides were used for real time PCR analysis (Fig.S7A):

- Ctail (recognises almost all isoforms): fw – GGTCCCATCATCAAGGTACG; rev – CATGGCTACCCTCGTTGTC
- SRR (recognises all isoforms): fw – ACTGAAGGAACAATGGACTCG; rev – CCAGAAAGAAGCAAAGCCTC
- PRR1 (recognises only PRR): fw – TCTACACCACTACCTACAGCA; rev – CAAGCCATCGCTACTATAGACG
- CH2 (recognises all isoforms): fw – GAAGTATCCCGTCCACGAG; rev – ACCACTCAATGTGCTCCTG
- CH2 (recognises only A*- and B*-type isoforms; Fig.1A): fw – CACCATCATCAGAGCTACCA; rev – CGTTCCATTGTTGCCACC

### Sequencing the *shot^V104^* breakpoint

The chromosomal breakpoint of *shot^V104^* was described to be in a 373bp region between bp73,398 and bp73,771 of the *shot* locus (Strumpf and Volk, 1998). We used an inverse PCR approach to determine the exact chromosomal break-point of *shot^V104^*. For this, genomic DNA of 200 homozygous *shot^V104^* embryos was isolated (Berkeley *Drosophila* Genome Project protocol; https://www.fruitfly.org/about/methods/inverse.pcr.html) and restricted with Sau96I. The restricted DNA was purified, diluted 10:1 and ligated into circular fragments. Using primer pairs designed to face towards the unknown region covering the breakpoint (fw: CCTGCTTTCAAACTAACATCCTGC; rev: CTGGCTGAATGGCAATTAAAGG), the circular DNA fragment containing the *shot^V10^*^4^ breakpoint region was amplified using a High Fidelity PCR Kit (Eppendorf and Roche). PCR products were gel-extracted, cloned into pDrive (Qiagen) and sequenced. The sequencing of one inverse PCR fragment showed a perfect alignment with wild-type genomic DNA until bp73.681 followed by an adenine and thymine-rich region. Using BLAST (https://flybase.org) we identified this region as part of the centromeric region of chromosome 2R (Fig.7). To confirm the breakpoint, sequence-specific primers were designed: a forward primer binding the wild-type *shot* region 100 bp upstream of the breakpoint (sense primer: TCTACGCTTGCGCTGCCCGCTCGCC) and three reverse primers; these were (1) antisense wt1 binding the wild-type region before the breakpoint: TTTGTACGCATTGGCATGGCAGATG; (2) antisense wt2 binding the wild-type region directly after: GGCAGATGCACAGATGCATTTATATACGC, and (3) antisense mutant 1 in the putative new *shot^V104^* sequence after the breakpoint: TGTTAGTTCTTATACAAGAAGATTCAATAAATAAAAGC. PCR results confirmed the breakpoint (Fig.7).

## Acknowledgements

Work reported here was made possible through the support of A.P. by the BBSRC (BB/C515998/1, BB/I002448/1, BB/M007553/1) and the Wellcome Trust (084561/Z/07/Z, 077748/Z/05/Z, 087820/Z/08/Z, 092403/Z/10/Z), of the BBSRC to N.S.S. (BB/R018960/1), and by parents as well as the Faculty of Life Sciences to Y.Q. The Manchester Bioimaging Facility microscopes used in this study were purchased with grants from the BBSRC, The Wellcome Trust and The University of Manchester Strategic Fund. The Fly Facility has been supported by funds from The University of Manchester and the Wellcome Trust (087742/Z/08/Z). Stocks obtained from the Bloomington *Drosophila* Stock Center (NIH P40OD018537) were used in this study.

---

**Fig.S1.**
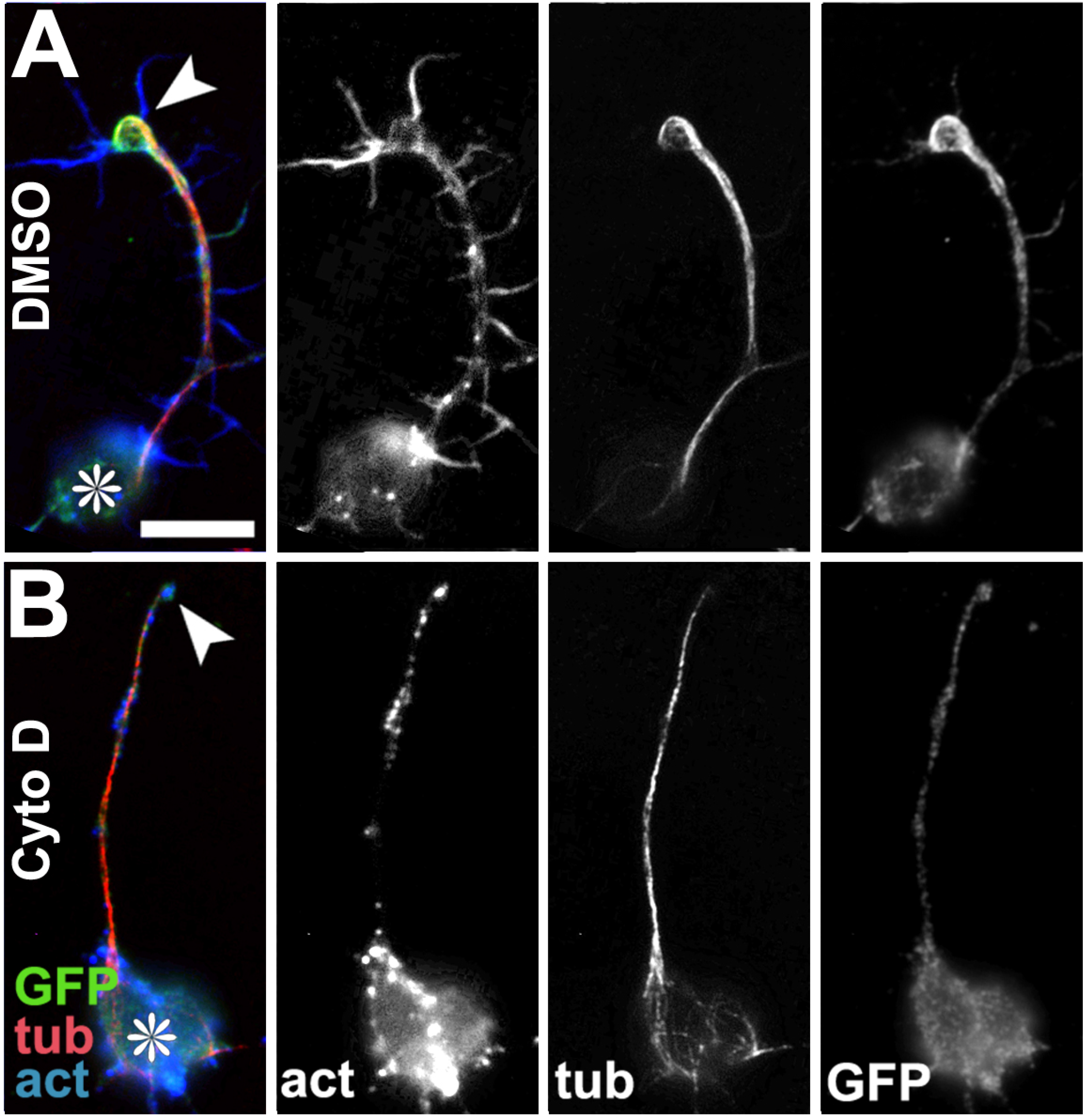
Impact of Cyto D on Shot-PE localisation. Primary neurons at 6-8 HIV on glass, treated with DMSO (control, **A**) or Cyto D (**B**) and stained for GFP (green), tubulin (red) and actin (blue); grayscale images on the right show single channels as indicated; asterisks indicate cell bodies, arrowheads the tip of axons; scale bar in A represents 10 µm in all images. For similar results compare Fig.6E,F.

**Fig.S2.**
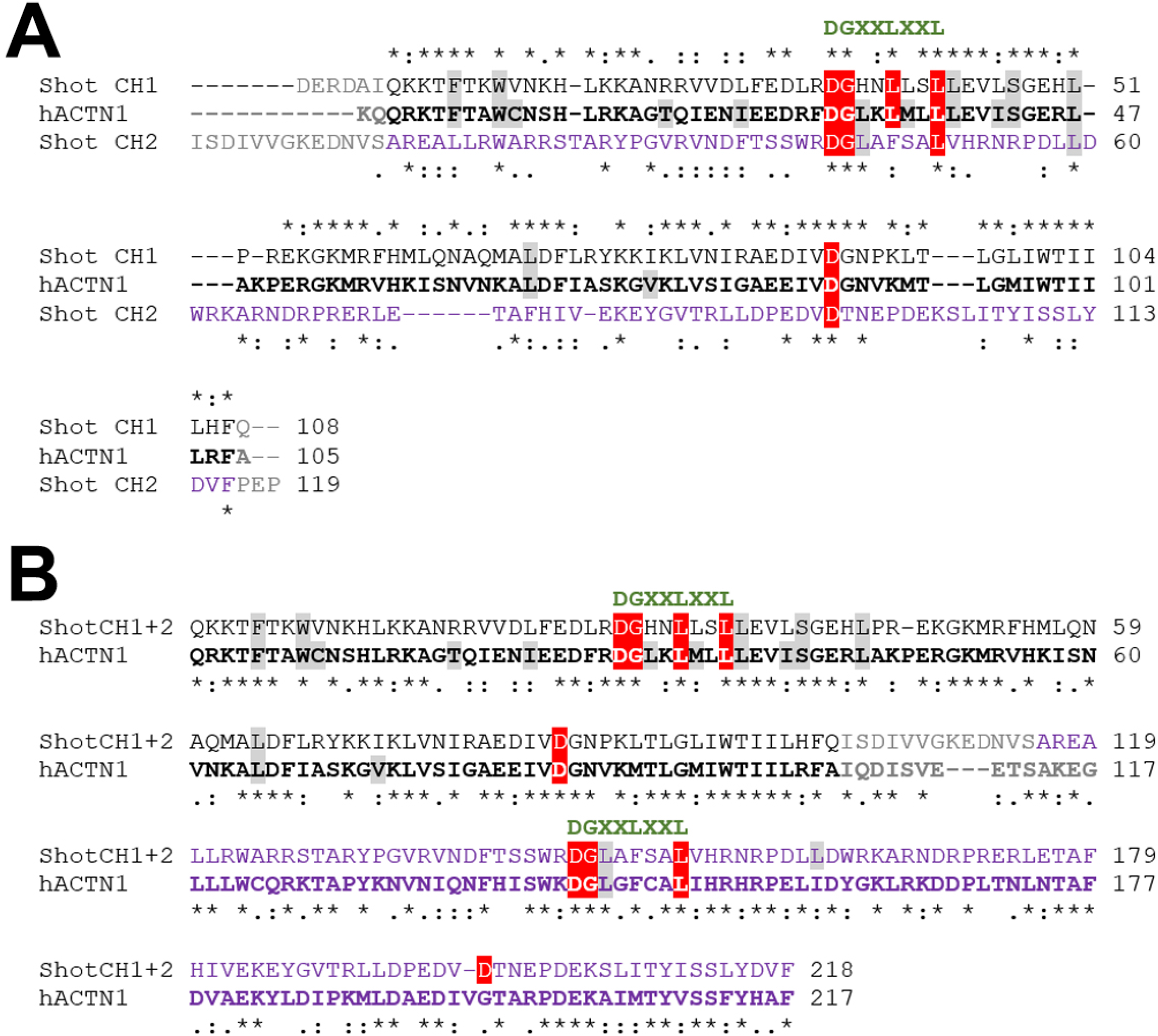
Comparison of the CH domains of Shot and human α1-actinin. Sequences are taken from the ACTN1-203 isoform (ensembl.org: [ENST00000394419.9](http://www.ensembl.org/Homo_sapiens/Transcript/Summary?db=core;g=ENSG00000072110;r=14:68874128-68979302;t=ENST00000394419)) and the Shot-PE isoform (flybase.org: FBtr0087618) and were aligned using UniProt Align (www.uniprot.org/align); asterisks indicate identical residues, dots and colons similar residues. **A**) Alignment of the first (CH1, black) and second (CH2, blue) CH domain of Shot with the first CH domain of hACTN1 indicates a high similarity of CH1 but considerable deviation of CH2 from the prototype CH domain; structurally important residues are colour-coded and the actin-binding consensus is provided above in green, as detailed elsewhere (Yin et al., 2020). **B**) Alignment of both CH domains as they occur in tandem in Shot and hACTN1 indicates a higher degree of identity and similarity of the two second CH domains (blue) to one another, than to the first CH domain (as shown for Shot CH2 in A).

**Fig.S3.**
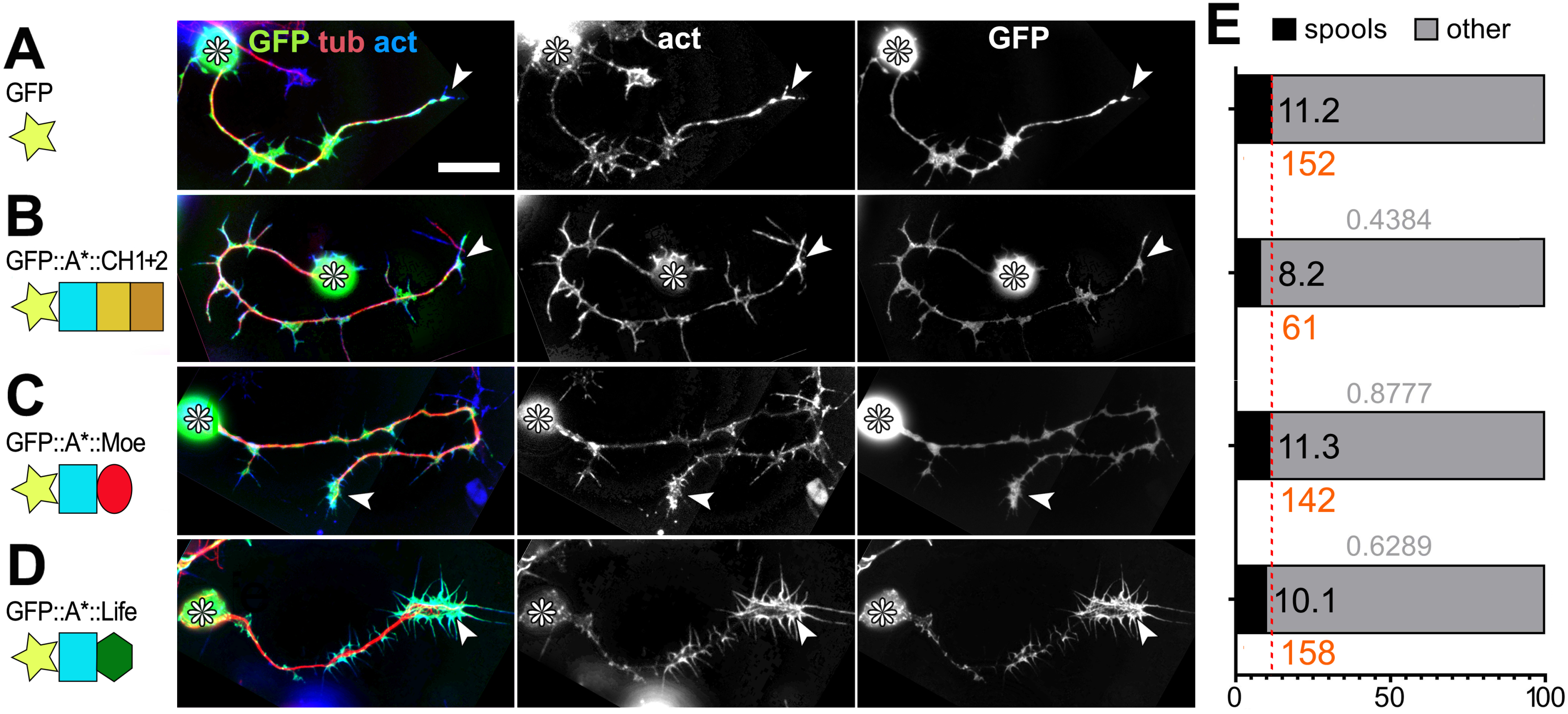
Only the Life-containing N-terminus shows strong F-actin association. **A-D**) Primary neurons at 24 HIV on ConA transfected with GFP (controls) or N-terminal constructs as indicated on the left (colour code as used in Fig.1); grayscale images on the right show single channels, as indicated; asterisks indicate cell bodies, arrowheads the tip of axons; scale bar in A represents 15 µm in all images. **E**) Bars correspond to experiments shown on the respective left and indicate the frequency of neurons with GCs that contain spools; number of neurons analysed are shown in orange, the percentage of neurons with GCs that contain spools in black in the bar, and P-values obtained via Chi^2^ tests on top of bars. Data were normalised to wild-type controls performed in parallel to all experiments (dashed red line).

**Fig.S4.** Animated GIF showing further examples of phenotypes induced by Shot-PE-Life::GFP expression. Primary neurons at 6-8 HIV on glass with *scabrous-Gal4*-induced expression of Shot-PE-Life::GFP, stained for tubulin (green), actin (red) and GFP (blue); the animation sequence shows single channels as grayscale images, as indicated top left in animation steps. Symbols indicate the following: asterisks, cell bodies; arrowheads, MT bundle split; arrows, ‘tennis racket’ spools; white curved arrows, unusual MT bundle malformations; open curved arrows, unusually bundled MTs in cell bodies. View or download: https://figshare.com/articles/figure/FigS4-Qu_al_gif/17056364.

**Fig.S5.**
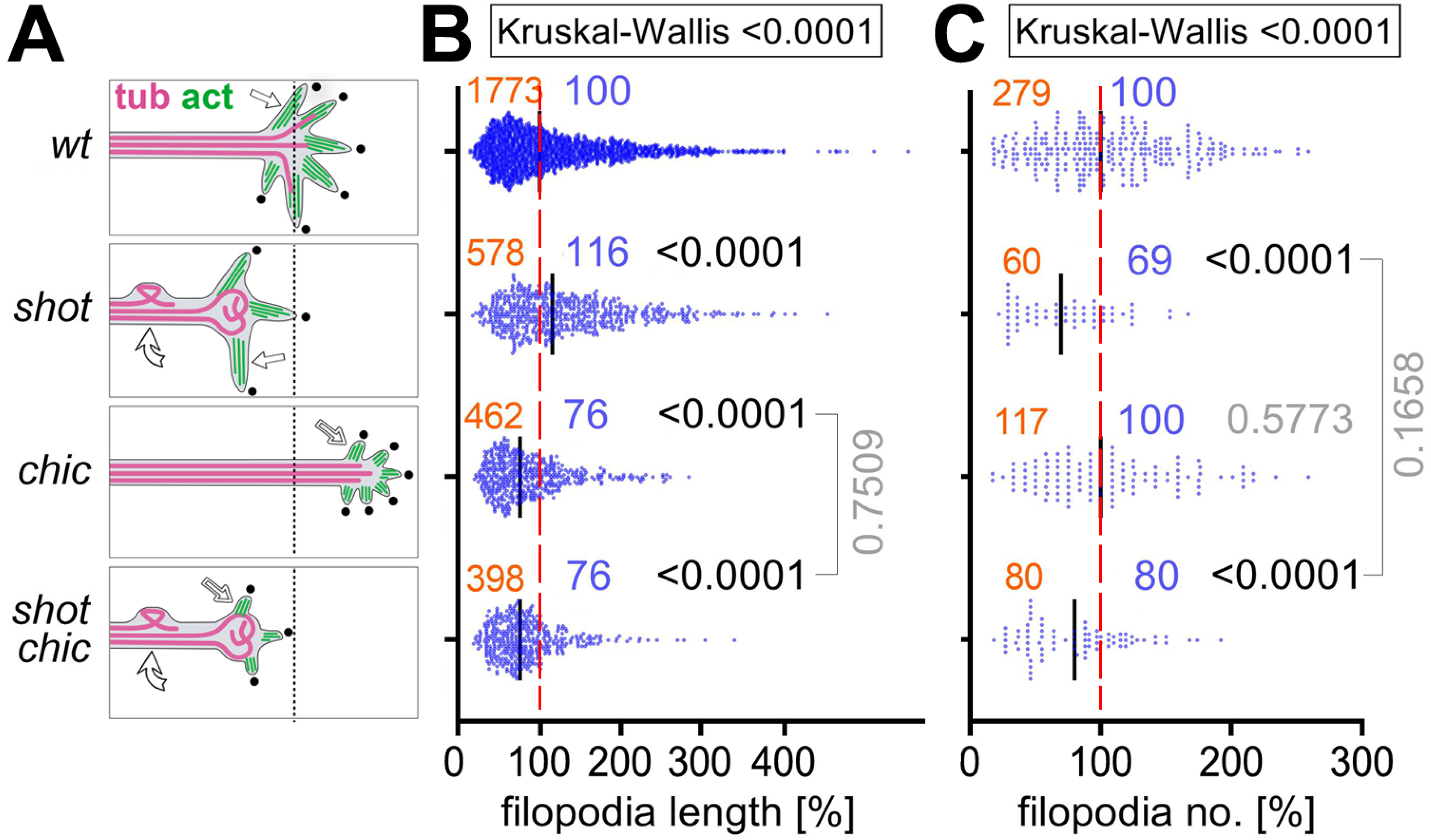
Filopodia data accompanying analyses shown in Fig.4B-E. **A)** Schematics on the left summarise the key findings for wild-type, *shot^3/3^*, *chic^221/221^* or *shot^3/3^ chic^221/221^* mutant primary neurons at 6-8 HIV on glass; indicated are: axon length (relative to black dashed line), filopodia number (black dots), filopodial length (white/open arrows) and MT disorganisation (curved arrows). **B,C**) Quantifications of filopodial lengths (B) and numbers of filopodia per neuron (C); numbers of analysed filopodia (B) or neurons (C) are indicated in orange; median values in blue, and P-values obtained via Mann–Whitney rank sum tests in black/grey; data were normalised to wild-type controls performed in parallel to all experiments (dashed red lines). Note that reduced filopodia numbers are due to regulatory roles of Shot in actin nucleation (Sánchez-Soriano et al., 2009), and shorter filopodia due to promoting roles of profilin/Chic in actin polymerisation (Gonçalves-Pimentel et al., 2011); both mechanisms are independent and therefore fully penetrant in the double-mutant constellation.

**Fig.S6.**
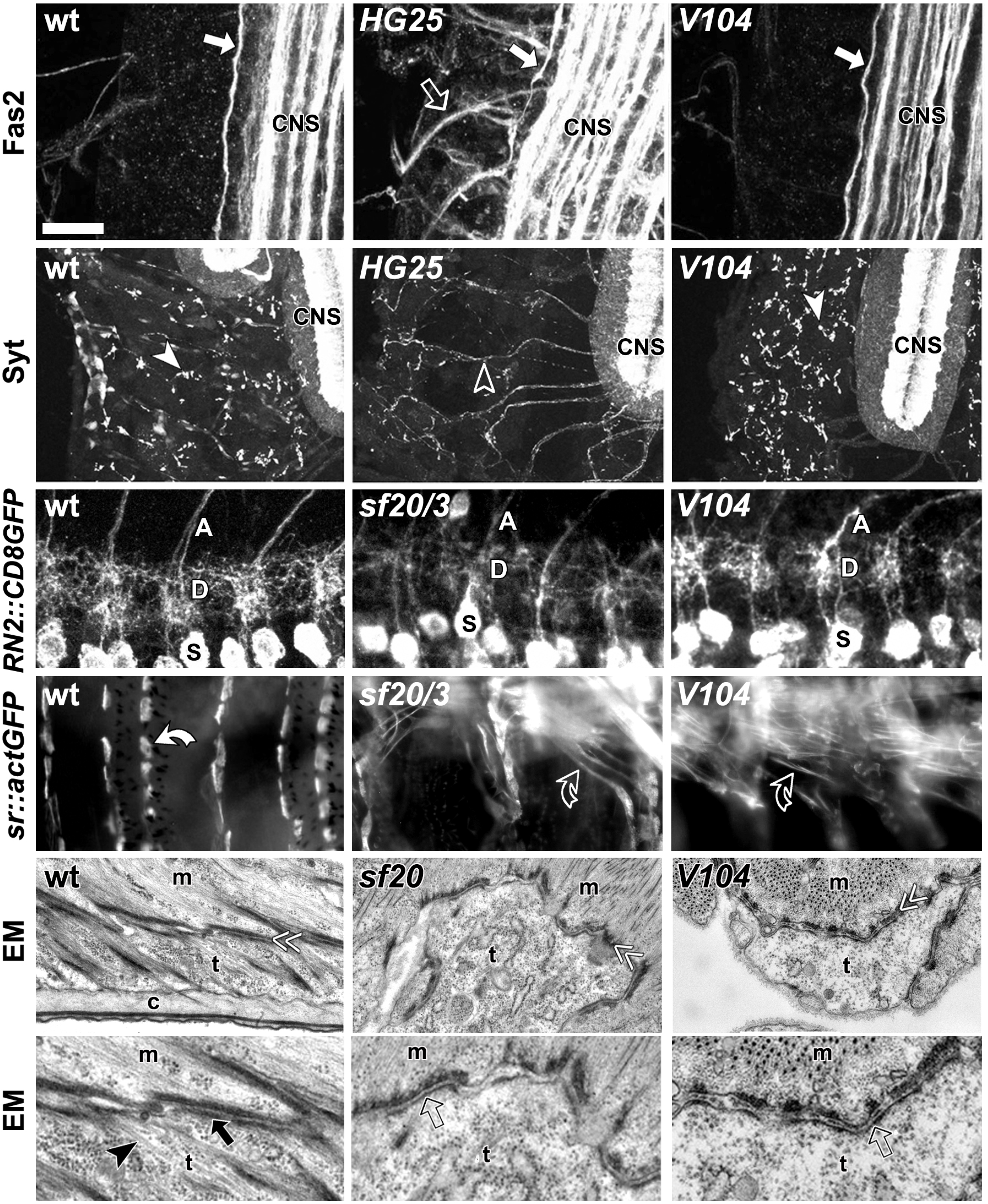
Phenotypes of *shot^V104^* mutant embryos. All images are taken from late stage 17 embryos of wild-type controls (wt; left), strong *shot* mutant alleles (HG25, sf20/3, sf20, middle; Lee et al., 2000; Prokop et al., 1998), and *shot^V104^* mutant embryos (right). **1^st^ row**: Fascilin 2-stained (Fas2) ventral nerve cords (part of the CNS; white arrow pointing at the most lateral longitudinal fascicle); only strong Shot deficiency causes the upregulation of Fas2 in nerve roots (open white arrow; Bottenberg et al., 2009; Prokop et al., 1998). **2^nd^ row**: flat dissected embryos stained for the synaptic marker Synaptotagmin (Syt; white arrowheads pointing at neuromuscular synapses); stained dots are severely reduced only by strong Shot deficiency (open arrowhead; Löhr et al., 2002). **3^rd^ row**: a detail of the ventral nerve cord expressing the membrane marker CD8::GFP driven by *eve^RN2^-Gal4* in a subset of motor neurons; somata (S), dendrites (D) and axons (A) are indicated: dendrites reduced only upon strong Shot deficiency (Bottenberg et al., 2009; Prokop et al., 1998). **4^th^ row**: flat dissected embryos expressing actin::GFP driven by *stripe-Gal4* (*sr*) in epidermal tendon cells (anchoring cells where muscles attach); only in the wild-type do tendon cells have their usual cell shapes (white curved arrow), whereas tendon cells become stretched (open curved arrows) upon strong Shot deficiency and in *shot^V104^*, indicating defects of the muscle-tendon junction (MTJ; Alves-Silva et al., 2008). **5^th^ & 6^th^ row**: micrographs of MTJs, central parts of which are shown twofold enlarged below; electron-dense MTJs (indicated by double chevrons) between muscles (m) and tendon cells (t) are properly formed in all genotypes, but the sub-membranous electron dense layer on the tendon cell side (black arrow in wt) is much thinner in the two *shot* mutant conditions (open arrows), and the characteristic MT arrays (arrowhead in wt) are absent, indicating the *shot*-specific tendon cell rupture phenotype (Prokop et al., 1998). This MTJ rupture in *shot^V104^* embryos is consistent with reports that Shot-PE-ΔCtail cannot rescue *shot* mutant tendon cell phenotypes (Alves-Silva et al., 2012), and that the Shot-PH isoform is not enriched in this cell type (Röper and Brown, 2003). Scale bar in A represents 20 µm in 1^st^, 40 µm in 2^nd^, 7 µm in 3^rd^, 30 µm in 4^th^, 1.2 µm in 5^th^ and 0.6 µm in 6^th^ row.

**Fig.S7.**
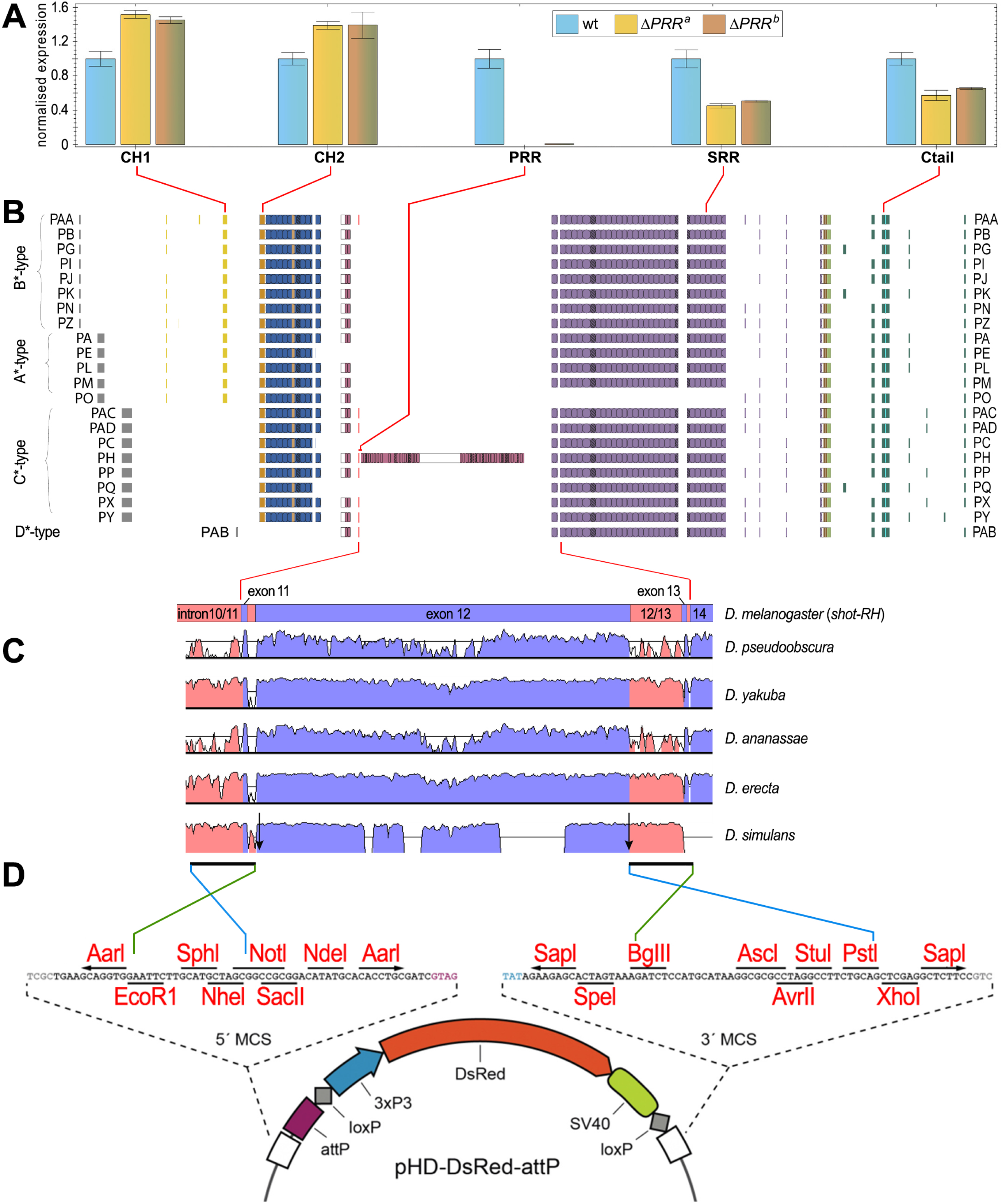
Generation and analysis of *shot^ΔPRR^*. **A**) Normalised data from quantitative RT-PCR analyses of embryos from wild-type and two independent CRISPR/Cas9 mutant lines (*shot^ΔPRRa^*, *shot^ΔPRRb^*) using probes against different exons (red lines) encoding different functional domains (indicated below graphs); they suggest that the PRR is deleted in the mutant strains, but that both versions of the allele cause severe expression changes of other exons suggestive of splice aberrations. **B**) A list of different splice variants of *shot* (modified from Voelzmann et al., 2017): names provided on the left and right, colour-coded as in Fig.1A and sorted by their A*-,B*-,C*- and D*-type N-termini (compare Fig.1A). **C**) Schematic representation of the *shot-RH* genomic sequence from exon 11 to 14 (position in the *shot* gene indicated by red lines) aligned with sequences from other *Drosophila* species (https://genome.lbl.gov/vista/customAlignment.shtml; introns in pink, exons in blue); the amplitude indicates the degree of evolutionary conservation, thus identifying areas that are not well-conserved and therefore suitable deletion sites less likely to affect important splice sites. Black arrows indicate the locations of the two guide RNAs for CRISPR/Cas9 incision (slightly removed from the 5’ end, and at the very 3’ end of exon 12), black bars the location of PCR-amplified flanking regions (covering part of intron 10/11 up to the end of intron 11/12; start of intron 12/13 to the start of exon 14) cloned into 5’ and 3’ multiple cloning sites (MCS) of *pHD-DsRed-attP* (D; vector scheme adapted from Gratz et al., 2014) using EcoRI/NotI and BglII/PstI restriction sites (for details see methods).

## Notes

https://figshare.com/articles/figure/FigS4-Qu_al_gif/17056364

### Competing Interest Statement

The authors have declared no competing interest.

## References

Alves-Silva, J., Hahn, I., Huber, O., Mende, M., Reissaus, A., Prokop, A. (2008). Prominent actin fibre arrays in Drosophila tendon cells represent architectural elements different from stress fibres. Mol. Biol. Cell 19, 4287–97 -- https://doi.org/10.1091/mbc.e08-02-0182

Alves-Silva, J., Sánchez-Soriano, N., Beaven, R., Klein, M., Parkin, J., Millard, T., Bellen, H., Venken, K. J. T., Ballestrem, C., Kammerer, R. A., Prokop, A. (2012). Spectraplakins promote microtubule-mediated axonal growth by functioning as structural microtubule-associated proteins and EB1-dependent +TIPs (Tip Interacting Proteins). J. Neurosci 32, 9143–58 -- https://doi.org/10.1523/JNEUROSCI.0416-12.2012

Amieva, M. R., Furthmayr, H. (1995). Subcellular localization of moesin in dynamic filopodia, retraction fibers, and other structures involved in substrate exploration, attachment, and cell-cell contacts. Exp Cell Res 219, 180–96 -- http://doi.org/10.1006/excr.1995.1218

Aumailley, M., Has, C., Tunggal, L., Bruckner-Tuderman, L. (2006). Molecular basis of inherited skin-blistering disorders, and therapeutic implications. Expert Rev Mol Med 8, 1–21 -- https://doi.org/10.1017/s1462399406000123

Bischof, J., Maeda, R. K., Hediger, M., Karch, F., Basler, K. (2007). An optimized transgenesis system for *Drosophila* using germ-line-specific varphiC31 integrases. Proc Natl Acad Sci U S A 104, 3312–7 -- https://doi.org/10.1073/pnas.0611511104

Blanchoin, L., Boujemaa-Paterski, R., Sykes, C., Plastino, J. (2014). Actin dynamics, architecture, and mechanics in cell motility. Physiol Rev 94, 235–63 -- http://www.ncbi.nlm.nih.gov/pubmed/24382887

Bottenberg, W., Sánchez-Soriano, N., Alves-Silva, J., Hahn, I., Mende, M., Prokop, A. (2009). Context-specific requirements of functional domains of the spectraplakin Short stop *in vivo*. Mech Dev 126, 489–502 -- https://doi.org/10.1016/j.mod.2009.04.004

Bray, D. (1984). Axonal growth in response to experimentally applied mechanical tension. Dev Biol 102, 379–89. -- https://doi.org/10.1016/0012-1606(84)90202-1

Buck, K. B., Zheng, J. Q. (2002). Growth cone turning induced by direct local modification of microtubule dynamics. J Neurosci 22, 9358–67 -- https://doi.org/10.1523/jneurosci.22-21-09358.2002

Budnik, V., Gorczyca, M., Prokop, A. (2006). Selected methods for the anatomical study of Drosophila embryonic and larval neuromuscular junctions. The fly neuromuscular junction: structure and function - Int. Rev. Neurobiol. 75, 323–74 -- https://doi.org/10.1016/s0074-7742(06)75015-2

Byers, T. J., Beggs, A. H., McNally, E. M., Kunkel, L. M. (1995). Novel actin crosslinker superfamily member identified by a two step degenerate PCR procedure. FEBS Lett 368, 500–4 -- https://doi.org/10.1016/0014-5793(95)00722-l

Campos-Ortega, J. A., Hartenstein, V. (1997). The embryonic development of Drosophila melanogaster. Springer Verlag, Berlin, pp. 227 -- https://www.springer.com/gb/book/9783662224915

Datar, A., Ameeramja, J., Bhat, A., Srivastava, R., Mishra, A., Bernal, R., Prost, J., Callan-Jones, A., Pullarkat, P. A. (2019). The roles of microtubules and membrane tension in axonal beading, retraction, and atrophy. Biophys J 117, 880–891 -- https://doi.org/10.1016/j.bpj.2019.07.046

Dent, E. W., Callaway, J. L., Szebenyi, G., Baas, P. W., Kalil, K. (1999). Reorganisation and movement of microtubules in axonal growth cones and developing interstitial branches. J Neurosci 19, 8894–908 -- https://doi.org/10.1523/JNEUROSCI.19-20-08894.1999

Dent, E. W., Gupton, S. L., Gertler, F. B. (2011). The growth cone cytoskeleton in axon outgrowth and guidance. Cold Spring Harb Perspect Biol 3, a001800 -- http://doi.org/10.1101/cshperspect.a001800

Dogterom, M., Koenderink, G. H. (2019). Actin–microtubule crosstalk in cell biology. Nat Rev Mol Cell Biol 20, 38–54 -- https://doi.org/10.1038/s41580-018-0067-1

Duchen, L. W., Strich, S. J., Falconer, D. S. (1964). Clinical and pathological studies of an hereditary neuropathy in mice (dystonia musculorum). Brain 87, 367–78 -- http://www.ncbi.nlm.nih.gov/pubmed/14188280

Edvardson, S., Cinnamon, Y., Jalas, C., Shaag, A., Maayan, C., Axelrod, F. B., Elpeleg, O. (2012). Hereditary sensory autonomic neuropathy caused by a mutation in dystonin. Ann Neurol 71, 569–72 -- http://www.ncbi.nlm.nih.gov/pubmed/22522446

Eyer, J., Cleveland, D. W., Wong, P. C., Peterson, A. C. (1998). Pathogenesis of two axonopathies does not require axonal neurofilaments. Nature 391, 584–7 -- https://doi.org/10.1038/35378

Franze, K., Janmey, P. A., Guck, J. (2013). Mechanics in neuronal development and repair. Annu Rev Biomed Eng 15, 227–51 -- http://www.ncbi.nlm.nih.gov/pubmed/23642242

Fritzsche, M., Lewalle, A., Duke, T., Kruse, K., Charras, G. (2013). Analysis of turnover dynamics of the submembranous actin cortex. Mol Biol Cell 24, 757–67 -- http://doi.org/10.1091/mbc.E12-06-0485

Fritzsche, M., Thorogate, R., Charras, G. (2014). Quantitative analysis of ezrin turnover dynamics in the actin cortex. Biophys J 106, 343–53 -- http://doi.org/10.1016/j.bpj.2013.11.4499

Fujioka, M., Emi-Sarker, Y., Yusibova, G. L., Goto, T., Jaynes, J. B. (1999). Analysis of an *even-skipped* rescue transgene reveals both composite and discrete neuronal and early blastoderm enhancers, and multi-stripe positioning by gap gene repressor gradients. Development 126, 2527–38 -- https://doi.org/10.1242/dev.126.11.2527

Geraldo, S., Khanzada, U. K., Parsons, M., Chilton, J. K., Gordon-Weeks, P. R. (2008). Targeting of the F-actin-binding protein drebrin by the microtubule plus-tip protein EB3 is required for neuritogenesis. Nat Cell Biol 10, 1181–9 -- https://doi.org/10.1038/ncb1778

Gonçalves-Pimentel, C., Gombos, R., Mihály, J., Sánchez-Soriano, N., Prokop, A. (2011). Dissecting regulatory networks of filopodia formation in a *Drosophila* growth cone model. PLoS ONE 6, e18340 -- https://doi.org/10.1371/journal.pone.0018340

Goriounov, D., Leung, C. L., Liem, R. K. (2003). Protein products of human Gas2-related genes on chromosomes 17 and 22 (hGAR17 and hGAR22) associate with both microfilaments and microtubules. J Cell Sci 116, 1045–58 -- https://doi.org/10.1242/jcs.00272

Goryunov, D., He, C. Z., Lin, C. S., Leung, C. L., Liem, R. K. (2010). Nervous-tissue-specific elimination of microtubule-actin crosslinking factor 1a results in multiple developmental defects in the mouse brain. Mol Cell Neurosci 44, 1–14 -- https://doi.org/10.1016/j.mcn.2010.01.010

Gratz, S. J., Ukken, F. P., Rubinstein, C. D., Thiede, G., Donohue, L. K., Cummings, A. M., O’Connor-Giles, K. M. (2014). Highly specific and efficient CRISPR/Cas9-catalyzed homology-directed repair in *Drosophila*. Genetics 196, 961–971 -- https://doi.org/10.1534/genetics.113.160713

Gregory, S. L., Brown, N. H. (1998). *kakapo*, a gene required for adhesion between cell layers in *Drosophila*, encodes a large cytoskeletal linker protein related to plectin and dystrophin. J. Cell Biol. 143, 1271–82 -- https://dx.doi.org/10.1083%2Fjcb.143.5.1271

Guha, S., Patil, A., Muralidharan, H., Baas, P. W. (2021). Microtubule sliding in neurons. Neurosci Lett, 135867 -- https://doi.org/10.1016/j.neulet.2021.135867

Hahn, I., Ronshaugen, M., Sánchez-Soriano, N., Prokop, A. (2016). Functional and genetic analysis of spectraplakins in *Drosophila*. Methods Enzymol 569, 373–405 -- https://doi.org/10.1016/bs.mie.2015.06.022

Hahn, I., Voelzmann, A., Liew, Y.-T., Costa-Gomes, B., Prokop, A. (2019). The model of local axon homeostasis - explaining the role and regulation of microtubule bundles in axon maintenance and pathology Neural Dev 14, 10.1186/s13064-019-0134-0 -- https://doi.org/10.1186/s13064-019-0134-0

Hahn, I., Voelzmann, A., Parkin, J., Fuelle, J. B., Slater, P. G., Lowery, L. A., Sanchez-Soriano, N., Prokop, A. (2021). Tau, XMAP215 and Eb co-operatively regulate microtubule polymerisation and bundle formation in axons. PLoS Genet 17, e1009647 -- https://doi.org/10.1371/journal.pgen.1009647

Halfon, M. S., Gisselbrecht, S., Lu, J., Estrada, B., Keshishian, H., Michelson, A. M. (2002). New fluorescent protein reporters for use with the *Drosophila* Gal4 expression system and for vital detection of balancer chromosomes. Genesis 34, 135–8 -- https://doi.org/10.1002/gene.10136

Harrison, R. G. (1910). The outgrowth of the nerve fiber as a mode of protoplasmic movement. J Exp Zool 9, 787–846 -- https://doi.org/10.1002/jez.1400090405

Hetrick, B., Han, Min S., Helgeson, Luke A., Nolen, Brad J. (2013). Small molecules CK-666 and CK-869 inhibit actin-related protein 2/3 complex by blocking an activating conformational change. Chem Biol 20, 701–12 -- https://doi.org/10.1016/j.chembiol.2013.03.019

Honnappa, S., Gouveia, S. M., Weisbrich, A., Damberger, F. F., Bhavesh, N. S., Jawhari, H., Grigoriev, I., van Rijssel, F. J., Buey, R. M., Lawera, A., Jelesarov, I., Winkler, F. K., Wuthrich, K., Akhmanova, A., Steinmetz, M. O. (2009). An EB1-binding motif acts as a microtubule tip localization signal. Cell 138, 366–76 -- https://doi.org/10.1016/j.cell.2009.04.065

Ka, M., Jung, E. M., Mueller, U., Kim, W. Y. (2014). MACF1 regulates the migration of pyramidal neurons via microtubule dynamics and GSK-3 signaling. Dev Biol 395, 4–18 -- http://www.ncbi.nlm.nih.gov/pubmed/25224226

Ka, M., Kim, W. Y. (2015). Microtubule-Actin Crosslinking Factor 1 is required for dendritic arborization and axon outgrowth in the developing brain. Mol Neurobiol -- http://doi.org/10.1007/s12035-015-9508-4

Kelso, R. J., Hudson, A. M., Cooley, L. (2002). *Drosophila* Kelch regulates actin organization via Src64-dependent tyrosine phosphorylation. J Cell Biol 156, 703–13 -- https://doi.org/10.1083/jcb.200110063

Kiehart, D. P., Galbraith, C. G., Edwards, K. A., Rickoll, W. L., Montague, R. A. (2000). Multiple forces contribute to cell sheet morphogenesis for dorsal closure in *Drosophila*. J Cell Biol 149, 471–90 -- https://doi.org/10.1083/jcb.149.2.471

Kodama, A., Karakesisoglou, I., Wong, E., Vaezi, A., Fuchs, E. (2003). ACF7: an essential integrator of microtubule dynamics. Cell 115, 343–354 -- https://doi.org/10.1016/s0092-8674(03)00813-4

Kolodziej, P. A., Jan, L. Y., Jan, Y. N. (1995). Mutations that affect the length, fasciculation, or ventral orientation of specific sensory axons in the *Drosophila* embryo. Neuron 15, 273–286 -- https://doi.org/10.1016/0896-6273(95)90033-0

Korenbaum, E., Rivero, F. (2002). Calponin homology domains at a glance. J Cell Sci 115, 3543–5 - - https://doi.org/10.1242/jcs.00003

Krieg, M., Stühmer, J., Cueva, J. G., Fetter, R., Spilker, K., Cremers, D., Shen, K., Dunn, A. R., Goodman, M. B. (2017). Genetic defects in β-spectrin and tau sensitize C. elegans axons to movement-induced damage via torque-tension coupling. Elife 6, e20172 -- https://doi.org/10.7554/eLife.20172

Kundu, T., Dutta, P., Nagar, D., Maiti, S., Ghose, A. (2021). Coupling of dynamic microtubules to F-actin by Fmn2 regulates chemotaxis of neuronal growth cones. J Cell Sci 134 -- https://doi.org/10.1242/jcs.252916

Lamoureux, P., Heidemann, S. R., Martzke, N. R., Miller, K. E. (2010). Growth and elongation within and along the axon. Dev Neurobiol 70, 135–49 -- http://doi.org/10.1002/dneu.20764

Lee, A. C., Suter, D. M. (2008). Quantitative analysis of microtubule dynamics during adhesion-mediated growth cone guidance. Dev Neurobiol 68, 1363–77 -- https://doi.org/10.1002/dneu.20662

Lee, S., Harris, K.-L., Whitington, P. M., Kolodziej, P. A. (2000). *short stop* is allelic to *kakapo*, and encodes rod-like cytoskeletal-associated proteins required for axon extension. J Neurosci 20, 1096–1108 -- https://doi.org/10.1523/jneurosci.20-03-01096.2000

Lee, S., Kolodziej, P. A. (2002). Short stop provides an essential link between F-actin and microtubules during axon extension. Development 129, 1195–1204 -- https://doi.org/10.1242/dev.129.5.1195

Lemieux, M. G., Janzen, D., Hwang, R., Roldan, J., Jarchum, I., Knecht, D. A. (2014). Visualization of the actin cytoskeleton: different F-actin-binding probes tell different stories. Cytoskeleton (Hoboken) 71, 157–69 -- https://doi.org/10.1002/cm.21160

Leterrier, C., Dubey, P., Roy, S. (2017). The nano-architecture of the axonal cytoskeleton. Nature Reviews Neuroscience 18, 713–726 -- http://dx.doi.org/10.1038/nrn.2017.129

Leung, C. L., Sun, D., Zheng, M., Knowles, D. R., Liem, R. K. H. (1999). Microtubule actin cross-linking factor (MACF): A hybrid of dystonin and dystrophin that can interact with the actin and microtubule cytoskeleton. J Cell Biol 147, 1275–1285 -- https://doi.org/10.1083/jcb.147.6.1275

Lin, C.-M., Li, P.-N., Chen, H.-J. (2021). Purification of subdomains of the plakin repeat domain in microtubule actin-crosslinking factor1b from bacteria. MC-Trans Biotechnol 12, e2 -- https://bio.mcu.edu.tw/sites/default/files/u3/file/2021/2021v12e2%20.pdf

Lin, C. M., Chen, H. J., Leung, C. L., Parry, D. A., Liem, R. K. (2005). Microtubule actin crosslinking factor 1b: a novel plakin that localizes to the Golgi complex. J Cell Sci 118, 3727–38 -- https://doi.org/10.1242/jcs.02510

Lindsley, D. L., Zimm, G. G. (1992). The genome of Drosophila melanogaster. Academic Press, pp. 1133 --

Littleton, J. T., Bellen, H. J., Perin, M. S. (1993). Expression of Synaptotagmin in *Drosophila* reveals transport and localization of synaptic vesicles to the synapse. Development 118, 1077–88 -- https://doi.org/10.1242/dev.118.4.1077

Löhr, R., Godenschwege, T., Buchner, E., Prokop, A. (2002). Compartmentalization of central neurons in *Drosophila*: a new strategy of mosaic analysis reveals localization of pre-synaptic sites to specific segments of neurites. J Neurosci 22, 10357–10367 -- https://doi.org/10.1523/JNEUROSCI.22-23-10357.2002

Lowery, L. A., van Vactor, D. (2009). The trip of the tip: understanding the growth cone machinery. Nat Rev Mol Cell Biol 10, 332–43 -- https://doi.org/10.1038/nrm2679

Lu, W., Lakonishok, M., Gelfand, V. I. (2015). Kinesin-1-powered microtubule sliding initiates axonal regeneration in *Drosophila* cultured neurons. Mol Biol Cell -- http://doi.org/10.1091/mbc.E14-10-1423

Luo, L., Liao, Y. J., Jan, L. Y., Jan, Y. N. (1994). Distinct morphogenetic functions of similar small GTPases: *Drosophila* Drac1 is involved in axonal outgrowth and myoblast fusion. Genes Dev. 8, 1787–1802 -- https://doi.org/10.1101/gad.8.15.1787

Marsick, B. M., San Miguel-Ruiz, J. E., Letourneau, P. C. (2012). Activation of ezrin/radixin/moesin mediates attractive growth cone guidance through regulation of growth cone actin and adhesion receptors. J Neurosci 32, 282–96 -- https://doi.org/10.1523/jneurosci.4794-11.2012

Millard, T. H., Martin, P. (2008). Dynamic analysis of filopodial interactions during the zippering phase of *Drosophila* dorsal closure. Development 135, 621–6 -- https://doi.org/10.1242/dev.014001

Miller, K. E., Sheetz, M. P. (2006). Direct evidence for coherent low velocity axonal transport of mitochondria. J Cell Biol 173, 373–81 -- http://doi.org/10.1083/jcb.200510097

Mlodzik, M., Baker, N. E., Rubin, G. M. (1990). Isolation and expression of *scabrous*, a gene regulating neurogenesis in *Drosophila*. Genes Dev 4, 1848–61 -- https://doi.org/10.1101/gad.4.11.1848

Mohan, R., John, A. (2015). Microtubule-associated proteins as direct crosslinkers of actin filaments and microtubules. IUBMB Life 67, 395–403 -- https://doi.org/10.1002/iub.1384

Prokop, A. (2020). Cytoskeletal organization of axons in vertebrates and invertebrates. J Cell Biol 219, e201912081 -- https://doi.org/10.1083/jcb.201912081

Prokop, A. (2021). A common theme for axonopathies? The dependency cycle of local axon homeostasis. Cytoskeleton 78, 52–63 -- https://doi.org/10.1002/cm.21657

Prokop, A., Beaven, R., Qu, Y., Sánchez-Soriano, N. (2013). Using fly genetics to dissect the cytoskeletal machinery of neurons during axonal growth and maintenance. J. Cell Sci. 126, 2331–41 -- http://dx.doi.org/10.1242/jcs.126912

Prokop, A., Küppers-Munther, B., Sánchez-Soriano, N. (2012). Using primary neuron cultures of *Drosophila* to analyse neuronal circuit formation and function. The making and un-making of neuronal circuits in Drosophila 69, 225–47 -- http://dx.doi.org/10.1007/978-1-61779-830-6_10

Prokop, A., Sánchez-Soriano, N., Gonçalves-Pimentel, C., Molnár, I., Kalmár, T., Mihály, J. (2011). DAAM family members leading a novel path into formin research. Commun Integr Biol 4, 538–42 -- http://doi.org/10.4161/cib.4.5.16511

Prokop, A., Uhler, J., Roote, J., Bate, M. C. (1998). The kakapo mutation affects terminal arborisation and central dendritic sprouting of Drosophila motorneurons. J Cell Biol 143, 1283–1294 -- https://doi.org/10.1083/jcb.143.5.1283

Qu, Y., Hahn, I., Lees, M., Parkin, J., Voelzmann, A., Dorey, K., Rathbone, A., Friel, C., Allan, V., Okenve Ramos, P., Sánchez-Soriano, N., Prokop, A. (2019). Efa6 protects axons and regulates their growth and branching by inhibiting microtubule polymerisation at the cortex. eLife 8, e50319 -- https://doi.org/10.7554/eLife.50319

Qu, Y., Hahn, I., Webb, S. E. D., Pearce, S. P., Prokop, A. (2017). Periodic actin structures in neuronal axons are required to maintain microtubules. Mol Biol Cell 28 296–308 -- https://doi.org/10.1091/mbc.e16-10-0727

Ramón y Cajal, S. (1890). À quelle époque apparaissent les expansions des cellules nerveuses de la moëlle épinière du poulet. Anatomischer Anzeiger 21-22, 609–639 --

Reuter, J. E., Nardine, T. M., Penton, A., Billuart, P., Scott, E. K., Usui, T., Uemura, T., Luo, L. (2003). A mosaic genetic screen for genes necessary for *Drosophila* mushroom body neuronal morphogenesis. Development 130, 1203–13 -- http://doi.org/10.1242/dev.00319

Riedl, J., Crevenna, A. H., Kessenbrock, K., Yu, J. H., Neukirchen, D., Bista, M., Bradke, F., Jenne, D., Holak, T. A., Werb, Z., Sixt, M., Wedlich-Söldner, R. (2008). Lifeact: a versatile marker to visualize F-actin. Nat Methods 5, 605–7 -- https://doi.org/10.1038/nmeth.1220

Roossien, D. H., Lamoureux, P., Van Vactor, D., Miller, K. E. (2013). *Drosophila* growth cones advance by forward translocation of the neuronal cytoskeletal meshwork in vivo. PLoS One 8, e80136 -- http://doi.org/10.1371/journal.pone.0080136

Röper, K., Brown, N. H. (2003). Maintaining epithelial integrity: a function for gigantic spectraplakin isoforms in adherens junctions. J Cell Biol 162, 1305–15 -- http://doi.org/10.1083/jcb.200307089

Sánchez-Soriano, N., Gonçalves-Pimentel, C., Beaven, R., Haessler, U., Ofner, L., Ballestrem, C., Prokop, A. (2010). *Drosophila* growth cones: a genetically tractable platform for the analysis of axonal growth dynamics. Dev Neurobiol 70, 58–71 -- https://doi.org/10.1002/dneu.20762

Sánchez-Soriano, N., Travis, M., Dajas-Bailador, F., Goncalves-Pimentel, C., Whitmarsh, A. J., Prokop, A. (2009). Mouse ACF7 and Drosophila Short stop modulate filopodia formation and microtubule organisation during neuronal growth. J Cell Sci 122, 2534–42 -- https://doi.org/10.1242/jcs.046268

Sanes, D. H., Reh, T. A., Harris, W. A., Landgraf, M. (2019). Development of the nervous system (4th edition). Academic Press, San Diego, pp. 360 -- https://www.elsevier.com/books/development-of-the-nervous-system/sanes/978-0-12-803996-0

Sjöblom, B., Ylänne, J., Djinovic-Carugo, K. (2008). Novel structural insights into F-actin-binding and novel functions of calponin homology domains. Curr Opin Struct Biol 18, 702–8 -- https://doi.org/10.1016/j.sbi.2008.10.003

Smith, D. H. (2009). Stretch growth of integrated axon tracts: extremes and exploitations. Prog Neurobiol 89, 231–9 -- http://doi.org/10.1016/j.pneurobio.2009.07.006

Strumpf, D., Volk, T. (1998). Kakapo, a novel *Drosophila* protein, is essential for the restricted localization of the neuregulin-like factor, Vein, at the muscle-tendon junctional site. J Cell Biol 143, 1259–1270 -- http://doi.org/10.1083/jcb.143.5.1259

Subramanian, A., Prokop, A., Yamamoto, M., Sugimura, K., Uemura, T., Betschinger, J., Knoblich, J. A., Volk, T. (2003). Short stop recruits EB1/APC1 and promotes microtubule assembly at the muscle-tendon junction. Curr Biol 13, 1086–95 -- https://doi.org/10.1016/s0960-9822(03)00416-0

Suter, D. M., Forscher, P. (2001). Transmission of growth cone traction force through apCAM-cytoskeletal linkages is regulated by Src family tyrosine kinase activity. J Cell Biol 155, 427–438 - - https://doi.org/10.1083/jcb.200107063

Tanaka, E., Sabry, J. (1995). Making the connection: cytoskeletal rearrangements during growth cone guidance. Cell 83, 171–6 -- http://www.ncbi.nlm.nih.gov/pubmed/7585934

Teng, J., Takei, Y., Harada, A., Nakata, T., Chen, J., Hirokawa, N. (2001). Synergistic effects of MAP2 and MAP1B knockout in neuronal migration, dendritic outgrowth, and microtubule organization. J Cell Biol 155, 65–76 -- https://doi.org/10.1083/jcb.200106025

Tessier-Lavigne, M., Goodman, C. S. (1996). The molecular biology of axon guidance. Science 274, 1123–33 -- https://doi.org/10.1126/science.274.5290.1123

Van Vactor, D. V., Sink, H., Fambrough, D., Tsoo, R., Goodman, C. S. (1993). Genes that control neuromuscular specificity in *Drosophila*. Cell 73, 1137–53 -- https://doi.org/10.1016/0092-8674(93)90643-5

Verheyen, E. M., Cooley, L. (1994). Profilin mutations disrupt multiple actin-dependent processes during *Drosophila* development. Development 120, 717–728 -- https://doi.org/10.1242/dev.120.4.717

Voelzmann, A., Liew, Y.-T., Qu, Y., Hahn, I., Melero, C., Sánchez-Soriano, N., Prokop, A. (2017). *Drosophila* Short stop as a paradigm for the role and regulation of spectraplakins. Semin Cell Dev Biol 69, 40–57 -- http://doi.org/10.1016/j.semcdb.2017.05.019

Voelzmann, A., Sánchez-Soriano, N. (2021). *Drosophila* primary neuronal cultures as a useful cellular model to study and image axonal transport Methods Mol Biol --

Warming, S., Costantino, N., Court, D. L., Jenkins, N. A., Copeland, N. G. (2005). Simple and highly efficient BAC recombineering using galK selection. Nucleic Acids Res 33, e36 -- https://doi.org/10.1093/nar/gni035

Winding, M., Kelliher, M. T., Lu, W., Wildonger, J., Gelfand, V. I. (2016). Role of kinesin-1-based microtubule sliding in *Drosophila* nervous system development. Proc Natl Acad Sci U S A 113, E4985–94 -- http://www.ncbi.nlm.nih.gov/pubmed/27512046

Woichansky, I., Beretta, C. A., Berns, N., Riechmann, V. (2016). Three mechanisms control E-cadherin localization to the zonula adherens. Nature Communications 7, 10834 -- https://doi.org/10.1038/ncomms10834

Xu, K., Zhong, G., Zhuang, X. (2013). Actin, Spectrin, and associated proteins form a periodic cytoskeletal structure in axons. Science 339, 452–6 -- http://doi.org/10.1126/science.1232251

Yin, L.-M., Schnoor, M., Jun, C.-D. (2020). Structural Characteristics, Binding Partners and Related Diseases of the Calponin Homology (CH) Domain. Front Cell Dev Biol 8 -- https://doi.org/10.3389/fcell.2020.00342

Zheng, J., Lamoureux, P., Santiago, V., Dennerll, T., Buxbaum, R. E., Heidemann, S. R. (1991). Tensile regulation of axonal elongation and initiation. J Neurosci 11, 1117–25 -- https://doi.org/10.1523/jneurosci.11-04-01117.1991

